# Discovery of novel drug combinations for oral squamous cell carcinoma through integrated analysis of single-cell RNA-sequencing datasets

**DOI:** 10.64898/2026.11.24.768808

**Authors:** Alexander Michael Grandits, Renate Wagner, Stefan Stoiber, Anna Sophie Berghoff, Agnes Forsthuber, Gerwin Heller, Lukas Kenner, Matthias Preusser, Thorsten Fuereder, Rotraud Wieser

## Abstract

Oral squamous cell carcinoma (OSCC) is associated with high morbidity and mortality, yet treatment options remain limited and resistance driven by intratumoral heterogeneity (ITH) is frequent. To characterize ITH and identify strategies to overcome it, we integrated twelve public single cell RNA sequencing datasets into an unprecedentedly large dataset and combined it with spatial transcriptomics. A novel, pathway-based clustering approach revealed seven highly recurrent and distinct tumor cell clusters, each defined by divergent cell-cell interactions, distinct positions along the cell trajectory tree, and spatially defined localization within tumors. To overcome ITH therapeutically, we sought to identify drug targets that were overexpressed in the vast majority of these highly diverse tumor cells, which was reasonably achieved only through pairwise target combinations. Pharmacological inhibition of LGALS1 plus CTSC, and LGALS1 plus TYMP, suppressed growth of OSCC spheroids, with further enhancement by cisplatin. Additional anti-tumor activity is expected *in vivo* through known effects of these inhibitors on the tumor microenvironment. Our work characterizes ITH in OSCC and reveals strategies to overcome it by broadly targeting malignant and microenvironmental drivers of OSCC, providing a basis for the rational design of novel combination therapies.

## 1 Introduction

Head and neck squamous cell carcinoma (HNSCC) is associated with high levels of pain, disfiguration, morbidity and mortality (1, 2). It mainly affects the oral cavity, oropharynx, hypopharynx and larynx, with some variation in clinical characteristics between these subsites (3). Etiological factors include chronic consumption of tobacco and/or alcohol, betel nut chewing, as well as human papillomavirus infection, the latter being predominantly involved in oropharyngeal carcinoma (4, 5). Even in the era of targeted therapies, standard-of-care treatment for newly diagnosed locally advanced HNSCC relies on surgery or platinum-based chemoradiation, which are used alone or in combination depending on the disease stage (3, 4). Molecularly guided therapies for HNSCC are restricted to the anti-epidermal growth factor receptor (EGFR) antibody cetuximab and immune checkpoint inhibitors like nivolumab and pembrolizumab (3).

The basis for the development of targeted therapies is the identification of recurrent genetic or transcriptional alterations, and these have been extensively catalogued for all major tumor entities in the past two decades. As examples, mutations in *TP53*, deletion of *CDKN2A*, or EGFR overexpression are frequently found in HNSCC (6). However, mutations or aberrant gene expression are not simply present in or absent from a patient’s tumor, but rather, their status varies between individual malignant cells. This phenomenon, termed intratumoral heterogeneity (ITH), is receiving growing attention as a key driver of therapy resistance and tumor recurrence, but has not yet been extensively investigated in HNSCC (7–9). Bulk transcriptomic studies have provided important insights regarding, e.g., gene dysregulation between tumor and corresponding normal tissue or the identification of prognostically relevant genes (10–12), but facilitate neither discrimination between the various cell types within a tumor nor the investigation of ITH. The advent of single-cell RNA-sequencing (scRNA-seq) methods has enabled these more in-depth analyses (13–15), and led to the publication of the first tumor cell atlas, established for melanoma, in 2016 (16). Because scRNA-seq dissociates cells from their spatial context, spatial transcriptomics (ST) was developed as a complementary technology that allows gene expression patterns to be aligned with histological images (17). As for other tumor types, a number of studies have applied these approaches to patients with HNSCC, resulting in novel insights, e.g., into the variety of cell types present in these tumors, their developmental relations, and their signaling interactions (reviewed in (18)).

The first scRNA-seq study published for HNSCC focused on the sub-entity, oral squamous cell carcinoma (OSCC) (19). It described the major cell types present in OSCC samples and reported a partial epithelial-mesenchymal transition (p-EMT) signature, which was, through immunohistochemistry, associated to the tumor leading edge (19). Re-analysis of these data with a focus on metabolic programs found that malignant cells exhibited upregulation of the largest number of metabolic pathways, but also cancer-associated fibroblasts were metabolically more active than myofibroblasts (20). An adaptation of scRNA-seq technology facilitating the simultaneous detection of human genes and those of the oral microbiota revealed that relevant bacterial genera were primarily associated with macrophages and with aneuploid epithelial cells highly expressing cancer progression-related signaling pathways (21). Several reports addressed the abundance and function of immune cells in HNSCC: cytotoxic, pre-dysfunctional, exhausted, and cycling CD8^+^ T cells, as well as regulatory CD4^+^ T cells, were found to be enriched in tumors compared to adjacent normal mucosa and peripheral blood (22–26). Poor responsiveness to neoadjuvant chemoimmunotherapy in HNSCC was associated with reduced levels of naïve T cells and memory B cells, as well as higher expression of immune-senescence related genes in lymphoid cells (27). In a consequently designed trial, the combination of ICI with senolytics yielded response rates superior to those of ICI monotherapy (27). ST was used, e.g., to characterize perineural invasion, a process associated with increased tumor aggressiveness, in OSCC. Nerves in close proximity to tumor tissue showed transcriptional responses to stress as well as degenerative processes, but also some indication of reprogramming to a survival mode (28). Combining scRNA-seq with ST, Su *et al.* uncovered molecular and cellular feedback mechanisms sustaining and enforcing the highly invasive properties of the OSCC leading edge (29). scRNA-seq and ST were also used to reveal novel potential drug targets in HNSCC (30–32).

Evidently, almost all of these studies also investigated the malignant cells themselves, and in several cases reported heterogeneous expression of tumor core and leading edge/EMT signatures (19, 29, 31). However, a more refined characterization of transcriptional ITH and of recurrent, tumor-relevant gene expression programs was hampered by the fact that malignant cells exhibited primarily copy number variation-driven, patient-specific gene expression patterns, which dominated their clustering and even remained largely conserved between primary tumors and metastases (19, 22, 30, 33, 34). Moreover, the high cost of scRNA-seq has limited most studies to small numbers of primary tumor samples, and especially in earlier studies, only modest numbers of epithelial cells were captured. To overcome these limitations, we integrated twelve publicly available scRNA-seq datasets comprising OSCC (n = 10) and normal oral mucosa (NOM; n = 2) samples, yielding more than 54,000 epithelial cells from the tumors of 68 patients with OSCC and from NOM of 23 healthy donors. We used this integrated scRNA-seq dataset to define recurrent tumor cell states based on pathway activity rather than expression of individual genes, and to identify drug targets that render the entire malignant cell population therapeutically vulnerable.

## 2 Results

### 2.1 Integration of publicly available scRNA-seq datasets from OSCC and normal oral mucosa

To establish an integrated scRNA-seq dataset containing large numbers of samples and of cells from OSCC and NOM, twelve scRNA-seq datasets were retrieved from the GEO database. Samples from sites other than the oral cavity and from diseases other than primary cancer were excluded, yielding 71 OSCC and 23 healthy donor samples (**Fig. 1A**, **Supplementary Tables S1** and **S2**) containing a total of 275,652 and 82,208 high-quality cells, respectively (**Supplementary Table S3**). The numbers of cells per sample, and of counts and genes per cell, varied between datasets, but not between the groups of OSCC and NOM samples (**Fig. 1B**, **Supplementary Fig. 1A**). Cells from the integrated dataset (hence termed the int_scRNA-seq dataset) clustered into the expected cell types (epithelial, endothelial, and skeletal muscle cells, fibroblasts, and various types of immune cells) in a dataset-of-origin independent manner (**Supplementary Fig. 1B-D**). Three OSCC samples did not contain any epithelial cells; in the remaining 68 specimens, a total of 49,093 epithelial cells were identified. The 23 NOM samples contained 5,460 epithelial cells. To further confirm the validity of the int_scRNA-seq dataset, protein coding genes significantly differentially expressed between epithelial cells from OSCC and NOM samples were compared between the int_scRNA-seq and TCGA datasets (**Fig. 1C**, **Supplementary Table S4A,B**). Of 7,682 and 7,153 differentially expressed genes, respectively, 3,332 were deregulated in the same direction in both datasets (*p* < 2.2×10^-16^, Pearson’s Χ^2^ test; Spearman’s ρ = 0.57), indicating consistent transcriptional regulation (**Fig. 1D**).

**Fig. 1:**
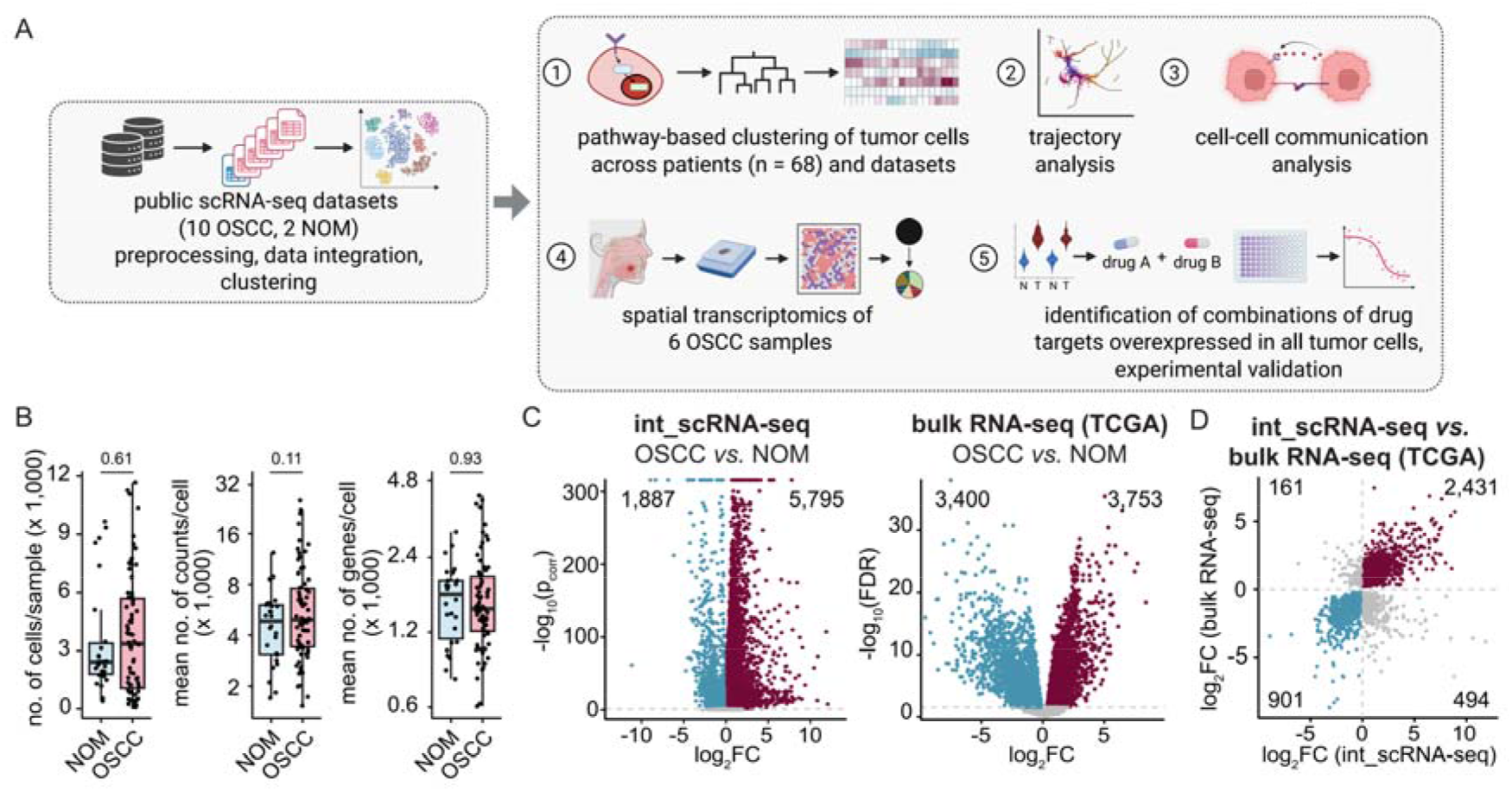
Integration and analysis of public scRNA-seq datasets from OSCC and normal oral mucosa. (A) Study overview. Ten scRNA-seq datasets containing a total of 71 OSCC samples and two scRNA-seq datasets containing a total of 23 normal oral mucosa (NOM) samples were retrieved from the Gene Expression Omnibus database, processed to yield an integrated scRNA-seq dataset, and analyzed as shown. Created in BioRender.com. Grandits, A. (2026) https://BioRender.com/skmndm9. (B) Total number of cells per sample, sample-level means of counts per cell, and sample-level means of detected genes per cell of NOM (n = 23) and OSCC (n = 71) samples. (C) Genes significantly (*p*_corr_ or FDR < 0.05) differentially expressed between tumor (n = 49,093) and NOM epithelial cells (n = 5,460) in the integrated scRNA-seq (int_scRNA-seq) dataset (left panel) and between OSCC (n = 268) and NOM (n = 19) samples from the TCGA dataset (right panel). (D) Comparison of the log_2_ fold changes of 3,987 genes that were significantly differentially expressed in both analyses. Spearman’s ρ = 0.57. FDR, false discovery rate; log_2_FC, log_2_ fold change; N, NOM; *p*_corr_, Bonferroni-corrected *p* value; T, OSCC.

In summary, integration of twelve publicly available scRNA-seq datasets yielded a combined int_scRNA-seq dataset allowing analyses of and comparisons between unprecedentedly large numbers of samples and cells from OSCC and NOM.

### 2.2 Identification of signaling states of tumor cells that are recurrent across patients and datasets

We decided to focus our first set of analyses of the int_scRNA-seq dataset on the tumor cells, with particular attention to ITH as a major driver of tumor evolution and therapy resistance (7). Aiming to discriminate tumor and normal cells among OSCC epithelial cells, CNV scores were first calculated for all epithelial cells. Notably, NOM epithelial cells, even though derived from donors without a diagnosis of OSCC, also appeared to bear CNVs according to this analysis (**Supplementary Fig. 2A**), and classification of normal and malignant cells was imperfect (area under the receiver operating characteristic curve = 0.84; **Supplementary Fig. 2B**). Nevertheless, in agreement with previous reports (22, 34, 35), most (95%) of the epithelial cells from OSCC samples would be called as malignant; we therefore considered it adequate to treat all epithelial cells from OSCC samples as tumor cells in further analyses.

Clustering of tumor cells using *Seurat* was driven by patient-specific CNVs, as reported previously (19, 22, 30, 33), but also by the number of reads per cell (which could reflect either biological or technical aspects; **Supplementary Fig. 3A,B**), and was thus deemed inadequate to identify relevant, recurrent tumor cell subpopulations. Therefore, instead of individual gene expression values, the activation status of the Hallmark Gene Sets (MSigDB), which characterize 50 fundamental biological states, was used to cluster tumor cells across patients and datasets through the *AUCell* algorithm. This analysis uncovered seven distinct clusters of tumor cells (termed AUCell 1 to AUCell 7; **Fig. 2A,B**, **Supplementary Fig. 4A**, **Supplementary Table S5**), with negligible effects of dataset-of-origin (**Supplementary Fig. 4B**). 40 samples contained all seven clusters, 12 only six, and in 16 samples, two or more clusters were missing (**Fig. 2C**). Samples in the latter groups contained 12- and 21-fold fewer tumor cells, respectively, than samples harboring all AUCell clusters (**Supplementary Fig. 4C**), indicating that the apparent absence of certain clusters in a subset of samples may reflect technical issues rather than biological differences.

**Fig. 2:**
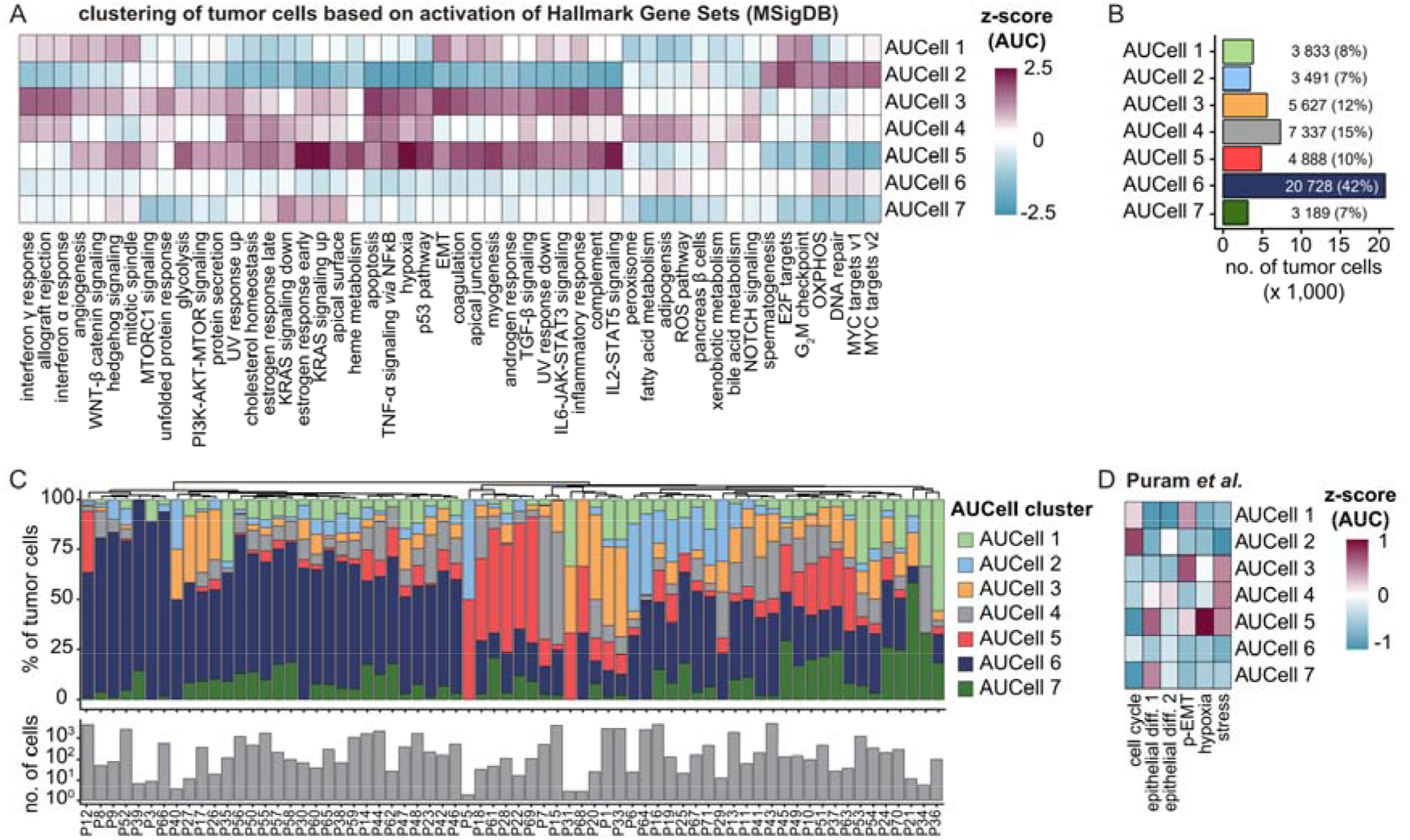
Identification of signaling states of tumor cells that are recurrent across patients and datasets. (A) Hierarchical clustering of tumor cells (n = 49,093) pooled from the OSCC samples (n = 68; another three samples did not contain any tumor cells) using the activation status of 50 Hallmark Gene Sets (MSigDB) and *AUCell*. (B) Total numbers and proportions of tumor cells in each AUCell cluster. (C) Top panel, relative proportions of the AUCell clusters in each sample. Samples were clustered to group those with similar compositions. Bottom panel, number of tumor cells per sample. (D) Enrichment of the gene sets defined by Puram *et al*. (19) in the AUCell clusters, determined through *AUCell*. AUC, area under the curve; diff., differentiation; MSigDB, Molecular Signatures Database; p-EMT, partial epithelial-mesenchymal transition.

AUCell 1 was strongly enriched for genes associated with epithelial-mesenchymal transition (EMT) and moderately for several proliferation-associated terms like E2F targets, mitotic spindle and G2M checkpoint. AUCell 2 showed more pronounced enrichment of proliferation-associated terms, along with upregulation of MYC target genes and oxidative phosphorylation (OXPHOS), but downregulation of many oncogenic signaling pathways, EMT, and apoptosis (**Fig. 2A**). In line with MYC acting as a general activator of transcription (36), AUCell2 was the cluster with the highest number of reads and genes per cell (**Supplementary Fig. 4D**). Moreover, consistent with their enrichment for cell cycle-related signatures, AUCell 1 and 2 contained smaller G_1_ and larger G_2_M fractions than the other clusters as well as NOM epithelial cells (**Supplementary Fig. 4E**). AUCell 3 exhibited upregulation of the inflammatory response and associated signaling pathways, as well as of apoptosis, P53 pathway, and EMT (**Fig. 2A**). AUCell 4 was characterized by high scores for TNF-α signaling *via* NF-κB, as well as for apoptosis and P53 pathway. AUCell 5 combined hypoxia, EMT, upregulation of glycolysis, downregulation of OXPHOS, and activation of multiple oncogenic signaling pathways with high scores for apoptosis and the P53 pathway, as well as downregulation of E2F and MYC target genes (**Fig. 2A**). Consistent with the latter observation, it contained the highest proportion of cells in G_1_ compared to all other clusters and to NOM cells (**Supplementary Fig. 4E**). AUCell 5 thus exhibited highly distinctive gene set activity patterns, with partially contradictory effects on tumor biology. AUCell 6 did not show strong enrichment or depletion of any of the Hallmark Gene Sets. AUCell 7 was mainly characterized by depletion of Hallmark Gene Sets, namely glycolysis, OXPHOS, reactive oxygen species, fatty acid metabolism, MYC targets, and E2F targets, and thus appears to contain cells with low metabolic and proliferative activity (**Fig. 2A**).

Enrichment of the tumor cell state signatures reported by Puram *et al*. (19) and Barkley *et al*. (37) in the AUCell clusters showed high concordance with the initial characterization (**Fig. 2D**, **Supplementary Fig. 5A**). Ingenuity Pathway Analysis (IPA) on genes significantly differentially expressed in one *vs*. all other clusters corroborated, among others, the positive association of AUCell 1 and 2, and the negative association of AUCell 5, with proliferation-related terms (**Supplementary Fig. 5B**; **Supplementary Table S6**). Similarly, telomere maintenance was activated in AUCell 2 and inhibited in AUCell 5 and 7. AUCell 2 was enriched, and AUCell 5 depleted, for OXPHOS, and in line with its hypoxic state, AUCell 5 was associated with increased VEGF signaling. IPA also revealed novel aspects of the AUCell clusters, e.g., the activation of keratinization, mostly reported as a good prognostic marker (38), and of the sheddase pathway, pivotal to the release of EGFR ligands from their membrane-bound precursors (39, 40), in AUCell 5 and 7 (**Supplementary Fig. 5B**; **Supplementary Table S6**). Analysis of upstream regulators confirmed MYC and E2F activity for AUCell 1 and 2 and TP53 activity for AUCell 3, 4, and 5 (**Supplementary Fig. 5C**).

Finally, a tumor core signature derived from ST of OSCC samples (31) was enriched in the hypoxic cluster AUCell 5 as well as in AUCell 7, with largely homogeneous upregulation of the respective markers. In contrast, the leading-edge markers were heterogeneously distributed in the remaining clusters (**Supplementary Fig. 5D**), suggesting that the leading edge is a more diverse region involving various tumor cell subpopulations.

In summary, our *AUCell*-based approach characterized transcriptional ITH in OSCC by identifying recurrent cell states distinguished by distinct pathway activity patterns. As examples, AUCell 2 was almost exclusively characterized by proliferation-associated features, while AUCell 5 combined reduced proliferation and activation of epithelial differentiation, P53 signaling and apoptosis with hypoxia, EMT and oncogenic signaling. These seemingly contradictory features align with literature reporting tumor core signatures as indicators of a good prognosis (31, 38), yet on the other hand describing hypoxia as a promoter of tumor stemness and therapy resistance (41, 42).

### 2.3 AUCell clusters are enriched in distinct branches of a cell trajectory tree and communicate through recurrent signaling pathways

Next, we interrogated the relationships between the AUCell clusters through cell trajectory and cell-cell communication analysis. Cell trajectory analysis yielded a tree with four branches (ending with clusters III, VI, VIII, and IX), in which tumor and NOM epithelial cells showed different distributions (**Fig. 3A**). AUCell 1, 3 and 5 were successively enriched along the branch ending with cluster VIII, and this branch was depleted of NOM epithelial cells (**Fig. 3B**). AUCell 2 was enriched in cluster IX, AUCell 4 in cluster VI, and AUCell 7 in cluster III, all of which represent the ends of distinct branches. NOM epithelial cells overlapped with AUCell 6 throughout the central parts of the tree, and with AUCell 4 and AUCell 7 at the ends of the branches terminating in clusters VI and III (**Fig. 3B**).

**Fig. 3:**
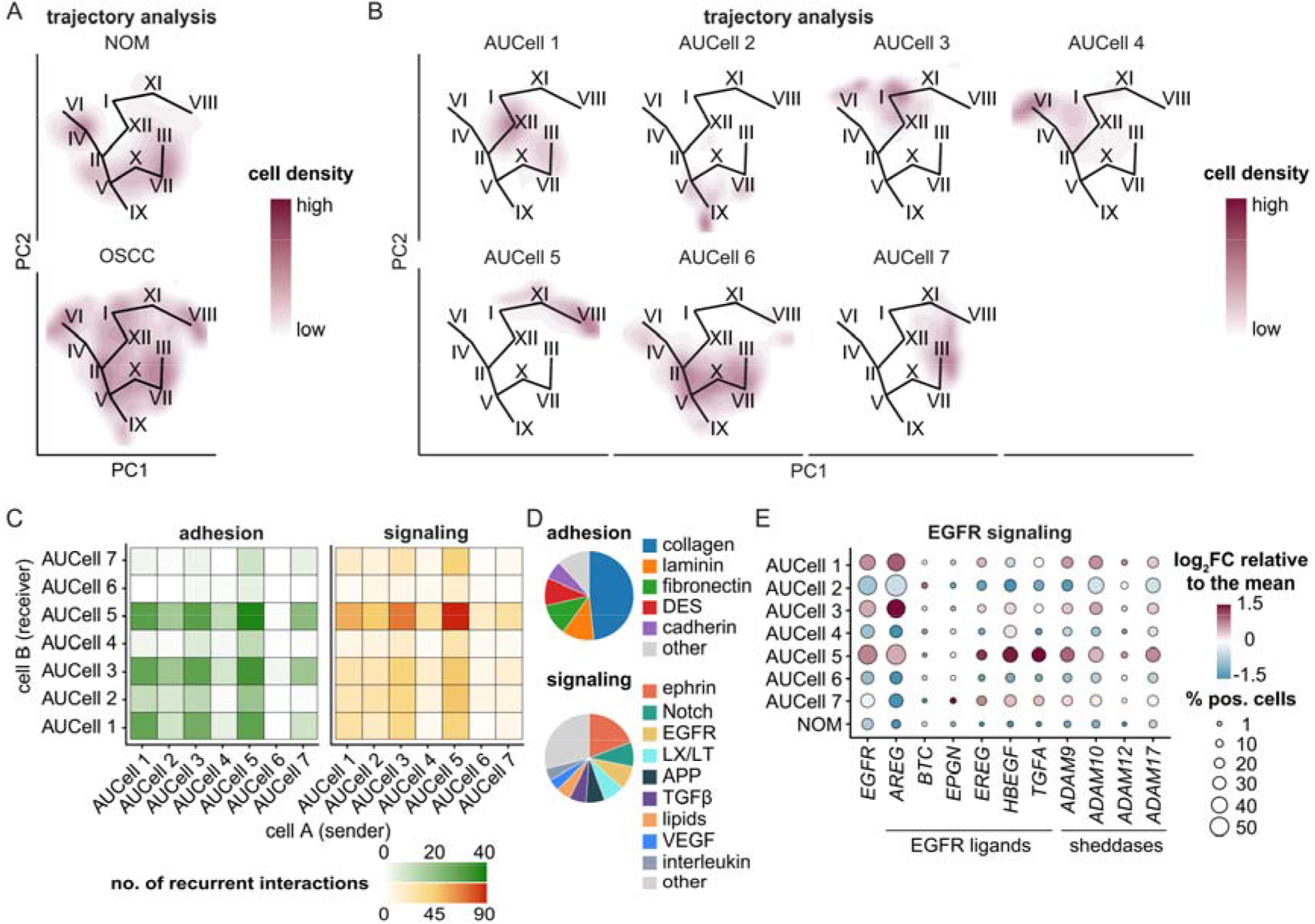
Cell trajectories of and cell-cell communication within and between AUCell clusters. (A) Cell trajectory analysis of tumor (n = 49,093) and normal oral mucosa (NOM) epithelial cells (n = 5,460), performed using *Lamian*. Top panel, NOM epithelial cells; bottom panel, tumor cells. (B) Distribution of the AUCell clusters along the trajectory tree. (C) Numbers of recurrent adhesion and signaling interactions within and between AUCell clusters. OSCC samples were analyzed individually using CellPhoneDB v5. Interactions that were found in >25% of the samples with ≥100 tumor cells and a mean of ≥1,000 detected genes per cell (n = 30) were considered recurrent. Classification of the interactions into “adhesion” and “signaling” was retrieved from CellPhoneDB v5. (D) Summary of the recurrent interactions for the classes “adhesion” and “signaling”. (E) Expression of *EGFR*, its ligands, and sheddases in the AUCell clusters and NOM epithelial cells. APP, amyloid precursor protein; DES, desmosome; log_2_FC, log_2_ fold change; LT, leukotriene; LX, lipoxin; NOM, normal oral mucosa; PC, principal component; pos., positive.

To identify recurrent, and therefore potentially tumor-relevant, cell-cell communication patterns within and between AUCell clusters, we first analyzed OSCC samples with ≥100 tumor cells (n = 42) individually using CellPhoneDB v5 (**Supplementary Fig. 6A**). The numbers of interactions per patient were highly variable and strongly correlated to the number of detected genes per cell (**Supplementary Fig. 6B,C**). Consequently, to avoid false-negatives due to low gene complexity, only samples with a mean of ≥1,000 detected genes per cell (n = 30) were included in the search for recurrent pathways, which we defined by their presence in >25% of patients. This criterion was met by 164 (47 adhesion, 104 signaling, 13 unclassified) of the 933 different interaction pathways that were found in any of the patients (**Supplementary Table S7**). The majority of these recurrent interactions (55% of the adhesion and 42% of the signaling interactions) involved AUCell 1, 3, and, most prominently, 5 (**Fig. 3C**), and many of them represent well known tumor promoting pathways, e.g., collagen-integrin interactions or EGFR signaling (43–45) (**Fig. 3D**). Interestingly, *EGFR* expression was strongest in AUCell 1, 3 and 5, and AUCell 5 additionally exhibited the highest proportions of cells with the highest expression levels of EGFR ligands and of the ADAM family proteases that convert them into their soluble forms (**Fig. 3E**).

In summary, these analyses show that AUCell 1, 3, and 5 are located on a separate branch of the cell trajectory tree that is largely depleted of NOM epithelial cells, and are most actively engaged in intratumoral cell-cell communication. AUCell 5 in particular forms the apex of this cell trajectory branch and exhibits the highest cell-cell communication activity, including oncogenic EGFR signaling.

### 2.4 AUCell clusters are associated with discrete pathological features in OSCC samples

To characterize the spatial distribution of the AUCell clusters and explore their potential associations with each other, non-malignant cell types, and/or pathological features, ST was performed on six OSCC samples (**Supplementary Table S8**). Necrotic, perinecrotic, vital tumor, and tumor-surrounding areas were identified based on HE staining (**Supplementary Fig. 7A**). Signals from Visium spots (n = 17,348) were deconvoluted into various non-epithelial cell types and the seven AUCell clusters using the smallest number of markers identified for any of the analyzed cell types (n = 71; AUCell 6) also for all other cell types to reduce detection bias. Still, the overall abundance of the AUCell clusters deviated partly from that in the scRNA-seq data (**Fig. 2B**, **4A**), which may be due to biological variation between samples (**Fig. 2C**) and/or to technical reasons like differences in gene detection technology or the mean fold-changes of the respective marker genes (**Supplementary Fig. 7B**). Nevertheless, all AUCell clusters, as well as the non-epithelial cell types typically present in the tumor microenvironment, were detected (**Supplementary Fig 7C,D**).

While most AUCell clusters appeared evenly distributed between vital tumor and perinecrotic areas, AUCell 5 was enriched in the perinecrotic areas in all patients (**Fig 4B**, **Supplementary Fig. 8A**). Confirming this observation, AUCell 5, and to a lesser extent AUCell 7, was associated with a lower density of cell nuclei, a characteristic of the perinecrotic area (**Fig. 4C**). Moreover, the tumor core signature, enriched in AUCell 5 and 7 in the int_scRNA-seq dataset (**Supplementary Fig. 5D**), was also enriched in the perinecrotic area (**Fig 4B**, **Supplementary Fig. 8B**).

**Fig. 4:**
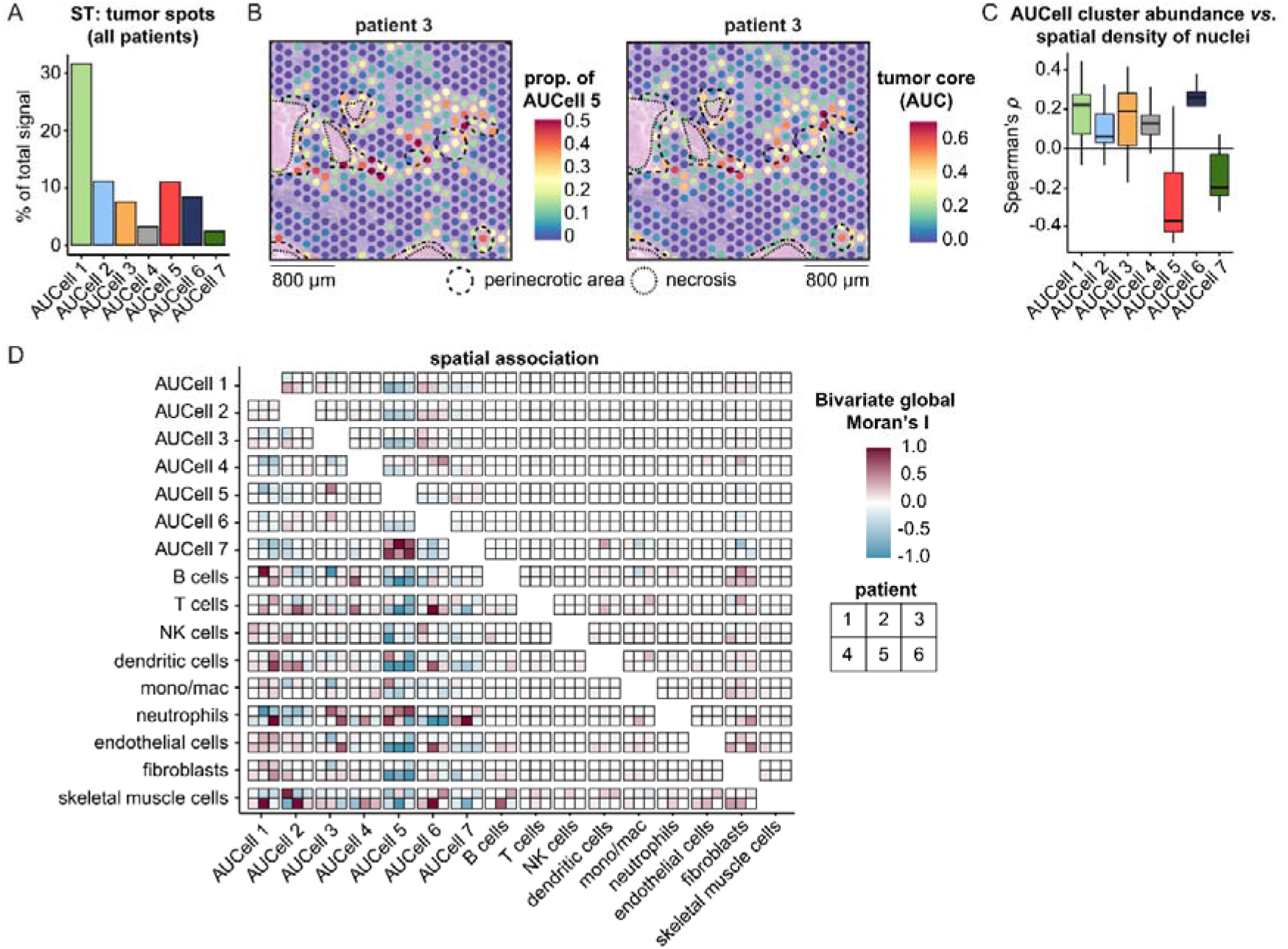
Spatial transcriptomics reveals associations of AUCell 5 with the perinecrotic region, distinct other cell types, and expression of EGFR signaling genes. (A) Proportion of AUCell clusters in tumor tissue-containing Visium spots (n = 5,167) from six OSCC samples subjected to spatial transcriptomics and spot deconvolution. (B) Representative tumor region of patient 3. Perinecrotic and necrotic areas are marked by dashed and dotted lines, respectively. Left panel, proportion of AUCell 5 signal as derived from spot deconvolution. Right panel, enrichment of the tumor core signature reported by Arora *et al*. (31), determined using *AUCell*. (C) Correlation of the proportion of AUCell clusters within tumor tissue-containing Visium spots with the spatial density of nuclei in their proximity. (D) Spatial associations between the AUCell clusters, and between AUCell clusters and other cell types in Visium spots containing tumor tissue was determined using the bivariate global Moran’s I. The analysis is not symmetrical because the proportion of one cell type in a given spot is associated with the spatially weighted mean proportion of another cell type in the neighboring spots. AUC, area under the curve; mono/mac, monocytes/macrophages; NK cells, natural killer cells; prop., proportion; ST, spatial transcriptomics.

To explore the spatial vicinity between cell types, the bivariate global Moran’s I (a direction-sensitive measure of spatial correlation) was used. While AUCell 1 and 6 were negatively associated to AUCell 7, and AUCell 5 to AUCell 1 and 2, the only positive association between tumor cell clusters occurred from AUCell 5 to AUCell 7 **Fig. 4D**). In line with the latter observation, and with the enrichment of AUCell 5 for the Hallmark Gene Set hypoxia (**Fig. 2A**), AUCell 5, and to a lesser extent AUCell 7, were negatively associated to endothelial cells. Additionally, AUCell 5 was negatively associated to fibroblasts, skeletal muscle cells, dendritic cells, B cells, T cells, and NK cells in the majority of samples (**Fig. 4D**).

In summary, ST analysis confirmed that AUCell 5 and 7, both enriched for the tumor core signature but otherwise exhibiting highly divergent pathway activity patterns, were enriched in the perinecrotic tumor core. Moreover, AUCell 5, whose signaling patterns reflect both the particular aggressiveness and relative indolence ascribed to tumor core cells, was negatively associated to several types of anti-tumor immune cells.

### 2.5 scRNA-seq data-driven discovery of novel combination treatments for OSCC

ITH, as exemplified by the AUCell clusters, provides tumors with a reservoir of cells with variable characteristics, some of which are bound to survive treatment and replenish the cells eradicated by therapy (7, 9). We therefore reasoned that the most valuable drug targets would be expressed at elevated levels in most or all tumor cells compared to NOM epithelial cells (**Fig. 5A**). The threshold for overexpression of a gene was defined as the 90^th^ percentile of its expression in NOM epithelial cells, and an OSCC sample was classified as potentially druggable if >90% of its tumor cells overexpressed one of 1,309 known drug targets from the IPA database. 15 of the 68 samples (22%) in the int_scRNA-seq dataset fulfilled this criterion for one or more genes. 13 genes were upregulated in one, seven in two, and one gene (*CTSC*) in four of the analyzed samples (**Supplementary Fig. 9A**). While this showed that in principle our approach was feasible, recurrent vulnerabilities were restricted to small proportions (3-6%) of patients. Therefore, all possible pairwise combinations of the 21 drug targets were interrogated whether one or both genes would be upregulated in >90% of a sample’s tumor cells. Seven combinations of six drug targets (*CTSC*, *LGALS1*, *LY6E*, *MIF*, *TNFRSF12A*, *TYMP*) met this criterion in a total of 32 samples (47%) that represented all ten original scRNA-seq datasets (**Fig. 5B**). Since potentially druggable samples had significantly more (1.7-fold) detected genes per cell than non-druggable samples (**Supplementary Fig. 9B**), increased assay sensitivity may further improve the identification of potentially treatable patients. Supporting the results of the scRNA-seq analyses, four drug targets were significantly upregulated in OSCC *vs*. NOM samples in all nine queried bulk gene expression datasets, and one each in eight and seven of them (**Fig. 5C**, **Supplementary Table S9**).

**Fig. 5:**
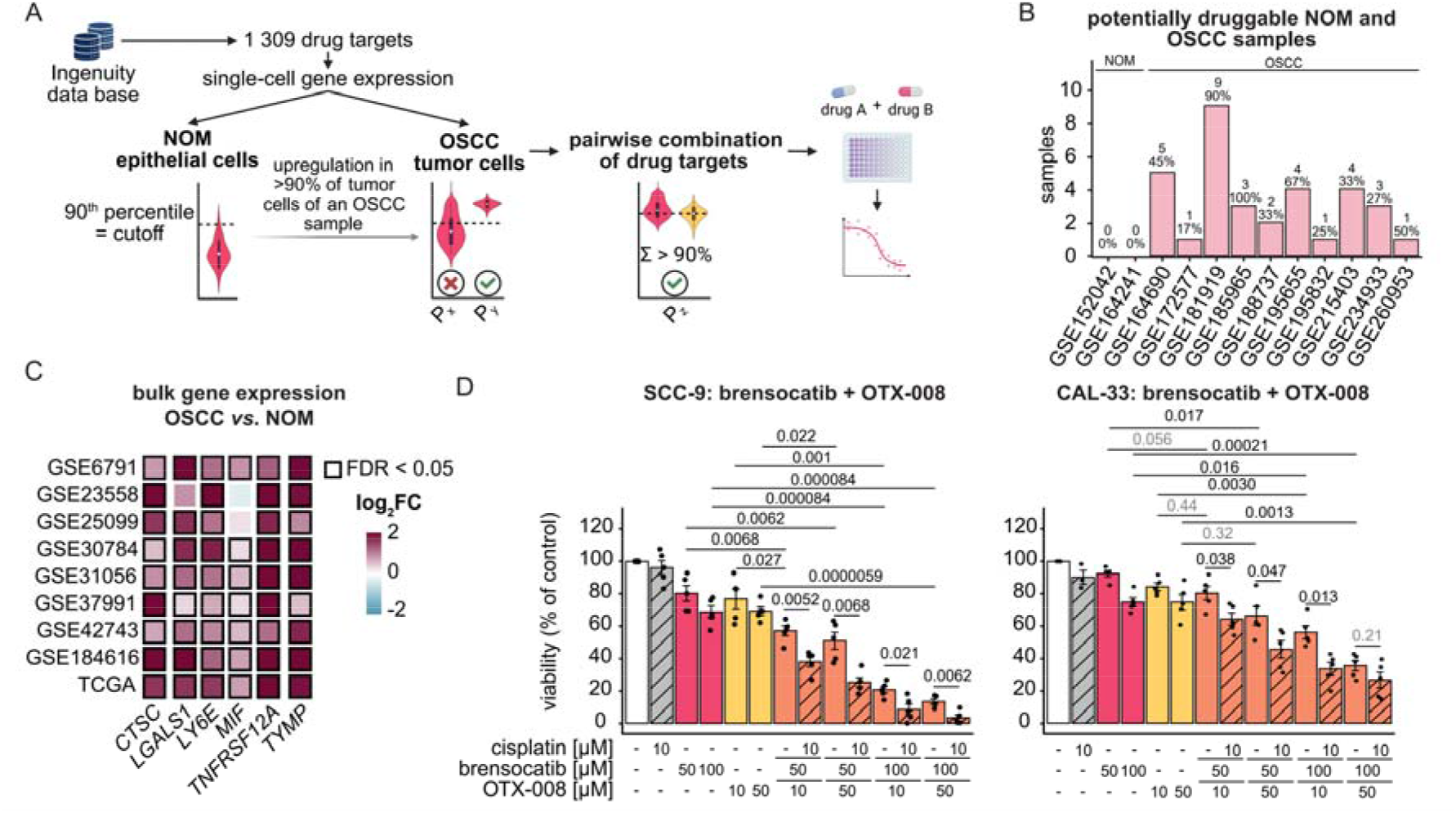
Combinations of targeted drugs identified through scRNA-seq exhibit synergistic effects on tumor spheroids that are further enhanced by cisplatin. (A) Schematic of the identification of patients potentially treatable with drug combinations based on the combined overexpression of the respective target genes in their tumor cells. Created in BioRender. Grandits, A. (2026) https://BioRender.com/ywv9plw. (B) Numbers and proportions of samples classified as druggable based on the expression of pairwise combinations of the target genes *CTSC*, *LGALS1*, *LY6E*, *MIF*, *TNFRSF12A* and *TYMP* in the NOM and OSCC datasets integrated into the int_scRNA-seq dataset. (C) Confirmation of the upregulation of the drug target genes in nine bulk gene expression datasets containing both OSCC and NOM samples. (D) 3D viability assay. Tumor spheroids of SCC-9 and CAL-33 cells were treated with the indicated concentrations of brensocatib, OTX-008 and/or cisplatin for 3 (CAL-33) or 5 (SCC-9) days and viability was quantified using the CellTiter-Glo® 3D Cell Viability Assay. Two-sided student’s *t* tests for predefined comparisons, followed by correction for multiple testing according to Bonferroni-Holm. Means ± SEM, n = 5. FDR, false discovery rate; log_2_FC, log_2_ fold change; NOM, normal oral mucosa.

To test the effectiveness of the drug combinations suggested by the *in silico* analyses, tumor spheroids of the OSCC cell lines SCC-9 and CAL-33, selected based on their expression of the target genes (**Supplementary Fig. 9C,D**) and their ability to form robust spheroids, were first treated with each drug individually. DLYE5953A (targeting LY6E) was not commercially available, and the monoclonal antibodies imalumab (targeting *MIF*) and enavatuzumab (targeting *TNFRSF12A*) showed very limited anti-proliferative effects as single agents (**Supplementary Fig. 10A**), thus were not followed up further. Brensocatib (targeting CTSC) plus OTX-008 (targeting LGALS1) had moderate single-agent activity (**Supplementary Fig. 10A**), but synergistic activity on spheroids from both cell lines (ZIP scores: 31.2 (SCC-9) and 10.9 (CAL-33); **Fig. 5D**, **Supplementary Fig. 10B**). OTX-008 plus tipiracil (targeting TYMP) showed synergistic and additive effects on SCC-9 (ZIP score: 27.0) and CAL-33 (ZIP score: 4.3) spheroids, respectively (**Supplementary Fig. 10C,D**). The combination of brensocatib with tipiracil had relatively modest overall activity (**Supplementary Fig. 10E,F**) and was not investigated further. Since chemotherapy is still a major pillar of OSCC therapy (3), we next added cisplatin to the two remaining drug combinations. Cisplatin significantly enhanced the effects of brensocatib plus OTX-008 and OTX-008 plus tipiracil on both CAL-33 and SCC-9 spheroids (**Fig. 5D, Supplementary Fig. 10C**). Together, these two drug combinations are predicted to be applicable to 14 (21%) of the 68 patients from the int_scRNA-seq dataset (**Supplementary Fig. 10G**). Importantly, all AUCell clusters showed similarly high expression of the drug targets and are therefore expected to be sensitive to treatment (**Supplementary Table S10**).

In summary, we identified novel drug combinations based on the expression of their respective targets in the vast majority of the tumor cells of a relevant proportion of OSCC patients. Two of these drug combinations exhibited synergistic activity against tumor spheroids from OSCC cell lines, and their effects were further enhanced by cisplatin. *In vivo,* additional therapeutic effects are expected because of the tumor-promoting effects of the selected drug target genes on stromal and immune cell populations (46–51). Together, these findings provide a basis for the further development of novel treatment strategies for OSCC.

## 3 Discussion

To characterize transcriptional ITH and to identify novel drug combinations for OSCC, we combined analyses of an scRNA-seq dataset, integrated from twelve public datasets and comprising large numbers of samples and cells from OSCC and NOM, with examinations of our own OSCC ST dataset. Identification of recurrent tumor cell states reflecting ITH through gene expression-based clustering was hampered by the strong impact of patient-specific CNVs (see also (19, 22, 30, 33)) and of the numbers of detected genes per cell. Therefore, we chose a novel approach and clustered tumors cells based on a pathway activation matrix rather than the gene expression matrix. This led to the identification of seven recurrent tumor cell clusters that were characterized by distinct pathway activation patterns, cell-cell communication profiles, cell trajectories, and spatial localizations within tumor tissue. Pursuing related goals on a pan-cancer level, Gavish *et al.* defined 41 meta-programs (MPs) in a cohort of 77 scRNA-seq datasets comprising 24 tumor types (52), of which OSCC was represented by a single dataset containing only ∼2,200 tumor cells (19). While theirs and our investigation concurred, e.g., regarding the proportions of proliferative (AUCell 1/2) and balanced (AUCell 6) OSCC cells, an in-depth comparison is limited by methodological differences: Gavish *et al*. assigned each cell to the single most active MP, while our analysis allowed simultaneous characterization of multiple activated pathways per cell, enabling the discovery of co-activation patterns such as EMT, E2F targets and several signaling pathways within the cells assigned to AUCell 1, or EMT, hypoxia, P53 signaling and apoptosis in AUCell 3 and 5. Of these clusters, AUCell 1 formed the root of a cell trajectory branch largely devoid of NOM epithelial cells. Along this branch, AUCell 1, 3, and 5 displayed progressive activation of EMT, hypoxia, and various pro-tumorigenic signaling pathways, as well as increased cell-cell interactions. On the other hand, tumor-suppressive pathways like apoptosis and P53 pathway were also activated in AUCell 3 and 5, and keratinization was upregulated in AUCell 5. AUCell 5 furthermore showed strong enrichment for a previously reported tumor core signature (31) and localized to the perinecrotic tumor core region in the ST analysis. The tumor core is an intensively researched region characterized by low vascularization, hypoxia, necrosis and keratinization (19, 31), and even though it has been associated with stemness, drug resistance and tumor aggressiveness, corresponding gene expression signatures predicted favorable patient outcomes (31, 38, 41, 42). This paradox mirrors the contradictory pathway activation patterns of AUCell 5. Surprisingly, AUCell 7, markedly different from AUCell 5 in pathway activation and cell-cell communication profiles, also exhibited upregulation of keratinization, was strongly enriched for the tumor core signature in the scRNA-seq data, and associated to features of the perinecrotic area in the ST analysis. Together, these findings highlight the functional complexity of the tumor core in OSCC, which manifests itself both on the inter- and intracellular level.

EGFR is considered a primary target of molecularly guided therapies for HNSCC, evidenced by the approval of cetuximab two decades ago (3, 4) and the recent development of several novel bispecific antibodies (53–55). However, although 80-90% of HNSCC samples are classified as EGFR-positive, only a subset of tumor cells expresses the receptor (56) (and **Fig. 3E**). We therefore hypothesized that targeting proteins overexpressed in the vast majority of tumor cells may expand the repertoire of potential treatment options for OSCC in a valuable manner, and analyzed the expression of a large set of druggable genes in our int_scRNA-seq dataset. Twenty-one target genes were overexpressed relative to NOM epithelial cells in >90% of the tumor cells of 1-4 samples, and seven pairwise combinations of six of these targets (*CTSC*, *LGALS1*, *LY6E*, *MIF*, *TNFRSF12A*, *TYMP*) classified almost half of the samples as potentially druggable, with 8-14 patients predicted to be sensitive to each combination. These numbers are likely underestimates, as non druggable samples exhibited markedly fewer detected genes per cell than druggable ones. Immunohistochemistry for target expression is expected to be more reproducible and potentially more sensitive than scRNA-seq, and is the method of choice in clinical settings.

Monoclonal antibodies against LY6E, MIF and TNFRSF12A, as well as the small molecule OTX-008 (targeting LGALS1) have demonstrated favorable toxicological profiles in preclinical studies or early-stage clinical trials (57–60). Tipiracil (targeting TYMP) in combination with trifluridine has already been approved for gastrointestinal cancer, and brensocatib (targeting CTSC) for non-cystic fibrosis bronchiectasis. Although monoclonal antibodies may exert direct anti-tumor activity *in vitro* (61), antibody-dependent cytotoxicity assays were considered more appropriate for evaluating their therapeutic potential. Consequently, the small molecule inhibitors brensocatib, OTX-008 and tipiracil were prioritized for *in vitro* testing. Tumor-promoting properties of CTSC, LGALS1 and TYMP have been demonstrated across multiple solid tumor models *in vitro* and *in vivo* (32, 35, 47, 48, 51, 62, 63). OTX-008 reduced the proliferation of OSCC cells *in vitro,* decreased tumor growth in OSCC and ovarian endometrioid adenocarcinoma xenograft models, and sensitized esophageal carcinoma cells to paclitaxel in *in vitro* and *in vivo* (60, 64, 65). Regarding inhibitors of CTSC and TYMP, to our knowledge our study is the first to investigate such agents in OSCC/HNSCC.

Of the three possible pairwise combinations, OTX-008 plus brensocatib and OTX-008 plus tipiracil synergistically reduced the viability of OSCC tumor spheroids, and their effects were further enhanced by cisplatin. Additional therapeutic effects of these drug combinations are expected *in vivo* because of the roles of their targets in tumor-promoting stromal and immune cell populations. Tumor-derived CTSC promoted neutrophil recruitment, induced neutrophil extracellular trap formation, and activated neutrophil serine proteases, collectively fostering a pro-tumorigenic inflammatory microenvironment and metastasis in breast cancer (46). In a mouse model of skin squamous cell carcinoma, CTSC expressed in fibroblasts and hematopoietic cells regulated immune cell infiltration and promoted angiogenesis and tumor growth (47). LGALS1 contributed to the accumulation of myeloid-derived suppressor cells, the conversion of myofibroblasts into cancer-associated fibroblasts, and elevated tumor vascular permeability (48–50). TYMP may foster tumor angiogenesis and T cell exhaustion (51). Overall, our approach suggests the triple combinations of OTX-008, brensocatib or tipiracil, and cisplatin as promising novel treatment strategies for at least a fifth of OSCC patients, targeting both the vast majority of tumor cells and pro-tumorigenic stromal and immune cells.

In summary, a novel pathway-based clustering strategy, applied to an unprecedentedly large OSCC scRNA-seq dataset and combined with ST analysis, identified seven highly recurrent tumor cell populations differing from each other by co-activation of specific sets of signaling pathways, cell-cell communication activity, and spatial localization, thus deepening our understanding of transcriptional ITH in OSCC. In addition, we implemented a straightforward approach for the discovery of novel drug combinations built on the premise that durable clinical responses require targeting of a maximal proportion of tumor cells. Based on scRNA-seq analyses and *in vitro* drug testing experiments, we propose two novel drug combinations that target the vast majority of tumor cells and concomitantly inhibit pro-tumorigenic stromal and immune components. This work lays the foundation for the development of novel therapeutic strategies capable of overcoming ITH by simultaneously and broadly targeting malignant and microenvironmental drivers of OSCC.

## 4 Methods

### 4.1 Retrieval of scRNA-seq datasets and data preprocessing and integration

To identify scRNA-seq datasets containing OSCC and/or NOM samples, the Gene Expression Omnibus (GEO; https://www.ncbi.nlm.nih.gov/geo/) was searched using the expression “((OSCC OR HNSCC OR “head and neck” OR oral) AND (“single cell” OR “single-cell” OR scRNA-seq OR scRNAseq) AND “Homo sapiens”[porgn: txid9606])”. As of May 2024, ten datasets containing OSCC samples and two datasets containing NOM samples were retrieved and included in the analysis (**Supplementary Table S1**). After exclusion of samples from sites other than the oral cavity and from diseases other than cancer, 71 OSCC and 23 NOM samples were available for analysis (**Supplementary Table S2**).

Unless otherwise stated, integration, processing and analysis of scRNA-seq data was performed in R v4.4.0 using the *Seurat* v5 package (66). To restrict the analyses to high quality cells, cells with >10% mitochondrial genes, <200 or >6,500 genes, >40,000 counts, or a novelty score (log_10_ ratio of the number of genes over the number of unique reads) of <0.8 were excluded. In addition, genes expressed in <0.1% of all cells were removed (**Supplementary Table S11**).

Samples were integrated using R*eciprocal Principal Component Analysis* that is implemented in *Seurat*. Cell types were assigned to the resulting clusters using *SingleR* and the cell type reference dataset *BlueprintEncodeData* from *celldex*. Epithelial cells that expressed *STATH* were classified as salivary gland cells (30) and excluded from further analysis. Three OSCC samples did not contain any epithelial cells, thus 68 samples remained for analyses of tumor cells.

Genes significantly differentially expressed (Bonferroni-corrected *p* value < 0.05) between epithelial cells from OSCC and NOM samples were identified using the *Seurat* function *FindMarkers*. Genes significantly (false discovery rate (FDR) < 0.05) differentially expressed between OSCC and NOM samples from the TCGA dataset were retrieved from the Xena platform using an adapted version of the Maayan lab’s Appyter bulk RNA-sequencing analysis pipeline (https://xenabrowser.net/). Both gene sets were filtered for protein coding genes.

### 4.2 InferCNV and receiver-operator curve analysis

Copy number variations (CNVs) were identified in epithelial cells from OSCC and NOM samples using *InferCNV* (19) and reference gene expression values derived from 55 000 randomly sampled non-epithelial cells. An overall CNV score was calculated for each cell using *InferCNV*-derived CNV classifications for each detectable gene (1: complete loss, 2: loss of one copy, 3: neutral, 4: addition of one copy, 5: addition of two copies, 6: addition of >2 copies) and the following equation:

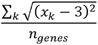

(x, CNV state; k, detected genes used by *InferCNV*, n_genes_ = number of detected genes) The receiver-operator curve and the corresponding area under the curve were calculated using *pROC*.

### 4.3 Pathway-based clustering of tumor cells and characterization of AUCell clusters

To cluster tumor cells (*i*.*e*., the epithelial cells from the OSCC samples, n = 49 093) based on the activation of the 50 Hallmark Gene Sets (Molecular Signatures Database, MSigDB https://www.gsea-msigdb.org/gsea/msigdb), the single-cell gene expression matrix was transformed into a pathway activation matrix using *AUCell*. Hierarchical clustering based on the Euclidean distance was performed using the base function *hclust*. The optimal number of clusters was selected based on the point at which additional clusters yielded diminishing gains in explained variance, as assessed using an elbow plot generated with the *factoextra* function *fviz_nbclust*.

Cell cycle phase was assigned to tumor (n = 49 093) and NOM epithelial cells (n = 5 473) using the *Seurat* function *CellCycleScoring*.

The activation of the gene sets reported by Puram *et al*. (19) and Barkley *et al*. (37) was compared between the AUCell clusters *via AUCell*. Ingenuity Pathway Analysis (IPA; QIAGEN, Venlo, Netherlands) was performed on the lists of genes significantly (Bonferroni-corrected *p* value < 0.05) differentially expressed in each AUCell cluster *vs*. all other AUCell clusters, established using the *Seurat* function *FindAllMarkers*. Pathways and upstream regulators with an FDR < 0.05 and a z-score >2 or <-2 were considered enriched or depleted, respectively, in a specific cluster.

Cell trajectory analysis of tumor and NOM epithelial cells was performed using *Lamian*. Cell-cell interactions within and between AUCell clusters were inferred for each sample with ≥100 tumor cells (n = 42) individually using CellPhoneDB v5 in python v3.12.7. Interactions with a score >25 were considered relevant. Interactions were considered recurrent if they were present in >25% of the analyzed OSCC samples with a mean of ≥1 000 detected genes per tumor cell (n = 30).

### 4.4 Spatial Transcriptomics

ST analysis of archival primary human OSCC samples was approved by the ethics committee of the Medical University of Vienna, Vienna, Austria (EK no. 1452/2022). Six treatment-naïve patients with OSCC were included; their characteristics are summarized in **Supplementary Table S8**. Sample processing, data acquisition, and primary data analysis were performed by the Core Facility Genomics, Medical University of Vienna, Vienna, Austria, according to the manufacturer’s recommendations. 5 µM sections from formalin-fixed paraffin-embedded samples were stained with HE and analyzed using the Visium v1 platform (10x Genomics, Pleasanton, CA, USA). Spaceranger v2.1.0 was used for preprocessing of the raw data, alignment of sequencing reads to the human genome (GRCh38), and generation of count matrices.

Based on the HE images, necrotic, perinecrotic, vital tumor, and tumor-surrounding areas were identified. Necrotic areas were excluded from gene expression analyses due to their very low read counts. The StarDist extension for QuPath v0.6.0 was used to obtain the positions of cell nuclei, which in turn were used to calculate their spatial density in a circular area (radius = 45 µm) around the center of each Visium spot.

Processing and analysis of ST data was performed in R v4.4.0. Signals from Visium spots were deconvoluted using *Robust Cell Type Decomposition* from *spacexr*. Marker genes for all represented cell types (*i*.*e*., the AUCell clusters as well as cells from the tumor microenvironment) were obtained from the int_scRNA-seq dataset using *FindAllMarkers*. The resulting gene lists were filtered for a Bonferroni-corrected *p* value < 0.05, a fold change > 1.5, and expression in >10% of the corresponding cells, resulting in 71 (AUCell 6) to 1 593 (endothelial cells) marker genes. To reduce detection bias, only the top 71 markers based on the Bonferroni-corrected *p* value were used for each cell type. Enrichment of the tumor core and leading edge signatures (31) in Visium spots was analyzed using *AUCell*.

To determine spatial associations between cell types, and between AUCell clusters and the expression of selected genes, the bivariate global Moran’s I was calculated between each tumor spot and the directly neighboring spots using the *spdep* function *moran_bv*.

### 4.5 Identification of drug target genes that are upregulated in tumor vs. NOM epithelial cells

1 309 genes listed as drug targets in the IPA database were analyzed for their upregulation in tumor *vs*. NOM epithelial cells. Since scRNA-seq gene expression data are right-skewed (97% of all genes are detected in less than 50% of the cells), the 90^th^ percentile of a gene’s expression in NOM epithelial cells (n = 5 473 cells, 23 samples) was defined as the lower threshold for upregulation. An OSCC sample (n = 68) was considered druggable if >90% of its tumor cells expressed a drug target gene above this threshold. Similarly, samples in which >90% of the tumor cells expressed one or both members of a pair of drug target genes above the respective threshold were classified as druggable by a combination therapy. Upregulation of the relevant drug target genes resulting from this analysis was confirmed in nine bulk RNA-sequencing datasets containing both OSCC and NOM samples. Paired samples were analyzed using paired tests.

### 4.6 Cell culture

The human OSCC cell lines CAL-27 (RRID:CVCL_1107), CAL-33 (RRID:CVCL_1108), SCC-4 (RRID:CVCL_1684), and SCC-25 (RRID:CVCL_1682) were acquired from the German Collection of Microorganisms and Cell Cultures GmbH (DSMZ, Braunschweig, Germany). SCC-9 cells (RRID:CVCL_1685) were kindly provided by Marcela Buchtová (Institute of Animal Physiology and Genetics, CAS, Brno, Czech Republic). Cell culture media are summarized in **Supplementary Table S12**. Cells were cultivated in a cell culture incubator at 37 °C, 5% CO_2_, and 95% relative humidity. Mycoplasma tests were performed regularly using the MycoAlert™ Mycoplasma Detection Kit (LT07-418, Lonza, Basel, Switzerland). To verify cell line identity, short tandem repeat (STR) profiling was performed by Eurofins Genomics (Luxembourg City, Luxembourg). STR profiles corresponded to profiles published by the DSMZ.

### 4.7 Quantification of drug target expression in OSCC cell lines

Preprocessed, log-transformed expression values of *CTSC*, *LGALS1*, *LY6E*, *MIF*, *TNFRSF12A*, and *TYMP* from 1 719 human cell lines were downloaded from the DepMap database (https://depmap.org/portal/, release data 26Q1).

Total RNA was isolated from OSCC cell lines using TRIzol® reagent (15596-018, Thermo Fisher Scientific) and reverse transcribed using the LunaScript™ RT SuperMix Kit (E3010L, NEB, Ipswich, MA, USA). For qRT-PCR, the GoTaq® qPCR master mix (A6001, Promega, Madison, WI, USA), custom-designed primers (**Supplementary Table S13**), and a StepOnePlus Real-Time PCR System (RRID:SCR_015805; Thermo Fisher Scientific) were used. All steps were performed according to the manufacturer’s standard protocols. In each biological replicate experiment, samples were measured in triplicate. Gene expression levels were normalized to those of *B2M* and to a reference sample using the ΔΔC_T_ method.

### 4.8 In vitro drug treatment of tumor spheroids

SCC-9 and CAL-33 cells were selected for *in vitro* testing due to their ability to grow in tumor spheroids. To generate tumor spheroids, subconfluent cells were trypsinized (25300-054, Thermo Fisher Scientific), and 3 000 cells were seeded per well of an ultra-low attachment plate (96-well format, U-bottom, 174925, Nunc A/S, Roskilde, Denmark) in keratinocyte SFM supplemented with 5 ng/mL EGF, 30 µg/mL pituitary extract (17005042, Thermo Fisher Scientific), and 5% fetal bovine serum (A5256701, Thermo Fisher Scientific), centrifuged for 15 min at 1 500 rpm, and incubated for 1 day. Then, tumor spheroids were treated with brensocatib (HY-101056), OTX-008 (HY-19756), tipiracil (HY-A0063A), enavatuzumab (HY-P99361), imalumab (HY-P99262; all from MedChemExpress LLC, Monmouth Junction, NJ, USA), and cisplatin (provided by the dispensary of the General Hospital, Vienna, Austria) at the concentrations shown in the figures for 5 (SCC-9) or 3 (CAL-33) days. Viability was measured using the CellTiter-Glo 3D Cell Viability Assay (G9683, Promega) according to the manufacturer’s instructions, and luminescence was read on a Varioskan LUX (RRID:SCR_026792; Thermo Fisher Scientific). Zip synergy scores were calculated in SynergyFinder 2.0 (http://synergyfinder.fimm.fi).

### 4.9 Statistics

*p* values, corrected *p* values and FDRs < 0.05 were considered statistically significant. Statistical tests were performed using GraphPad Prism v6 (GraphPad Software, San Diego, CA, USA) or R v4.4.0. The two-sided Student’s *t*-test and one-way ANOVA followed by Bonferroni’s post-hoc test were used to compare means of two or multiple groups, respectively. The two-way ANOVA followed by Bonferroni’s post-hoc test was used for multiple groups with two or more factors. Pearson’s Χ² test was used to test for relationships between categorical variables.

Three to five independent biological replicates were performed for qRT-PCR and drug treatment experiments. For the latter, a pre-defined set of two-sided student’s *t*-tests was performed and *p* values were corrected using the Bonferroni-Holm method. Corrected *p* values of all tested pairs are shown in the panels. Data are presented as means ± SEM (standard error of the mean) of the mean values derived from technical replicates.

## List of abbreviations

CNV: copy number variation
DSMZ: Deutsche Sammlung von Mikroorganismen und Zellkulturen (German Collection of Microorganisms and Cell Cultures GmbH)
EGFR: epidermal growth factor receptor
EMT: epithelial-mesenchymal transition
FDR: false discovery rate
GEO: Gene Expression Omnibus
HNSCC: head and neck squamous cell carcinoma
int_scRNA-seq dataset: integrated scRNA-seq dataset
IPA: Ingenuity Pathway Analysis
ITH: intratumoral heterogeneity
MSigDB: Molecular Signatures Database
MP: meta-program
NOM: normal oral mucosa
OXPHOS: oxidative phosphorylation
OSCC: oral squamous cell carcinoma
p-EMT: partial epithelial-mesenchymal transition
scRNA-seq: single-cell RNA-sequencing
SEM: standard error of the mean
ST: spatial transcriptomics
STR: short tandem repeats
TCGA: The Cancer Genome Atlas

## Declarations

### Ethics approval

ST analysis of archival primary human OSCC samples was approved by the ethics committee of the Medical University of Vienna, Vienna, Austria (EK no. 1452/2022).

### Consent for publication

Not applicable.

### Availability of data

The scRNA-seq data analyzed in this study are available in the GEO at <u>GSE152042</u>, <u>GSE164241</u>, <u>GSE164690</u>, <u>GSE172577</u>, <u>GSE181919</u>, <u>GSE185965</u>, <u>GSE188737</u>, <u>GSE195655</u>, <u>GSE195832</u>, <u>GSE215403</u>, <u>GSE234933</u> and <u>GSE260953</u>. Spatial transcriptomics data were deposited in the GEO (accession number <u>GSE325160</u>).

### Authors’ contributions

Conceptualization, AMG, RW; methodology, AMG, RWa, ASB, AF, GH, RW; validation, AMG, RW; formal analysis, AMG; investigation, AMG, RWa, SS, RW; writing-original draft preparation, AMG, RW; writing-review and editing: all authors; visualization, AMG; provision of patient samples, LK; supervision, MP, TF, RW; project administration, RW; funding acquisition, MP, RW.

All authors have read and agreed to the published version of the manuscript.

## Acknowledgements

This research was funded in whole or in part by the Austrian Science Fund (FWF) [grant 10.55776/PIN1127624]. For open access purposes, the author has applied a CC BY public copyright license to any author accepted manuscript version arising from this submission. Funding for this work was provided by the institution (Division of Oncology, Department of Medicine I, Medical University of Vienna).

The authors thank Catarina de Almeida Diogo de Bettencourt e Ávila, Martin Bilban, and Sophia Derdak from the Core Facilities of the Medical University of Vienna for providing spatial transcriptomics services and primary data analysis.

Large language models were used for language editing and debugging of R codes.

