## Supplementary Figures for "Discovery of novel drug combinations for oral squamous cell carcinoma through integrated analysis of single-cell RNA-sequencing datasets"

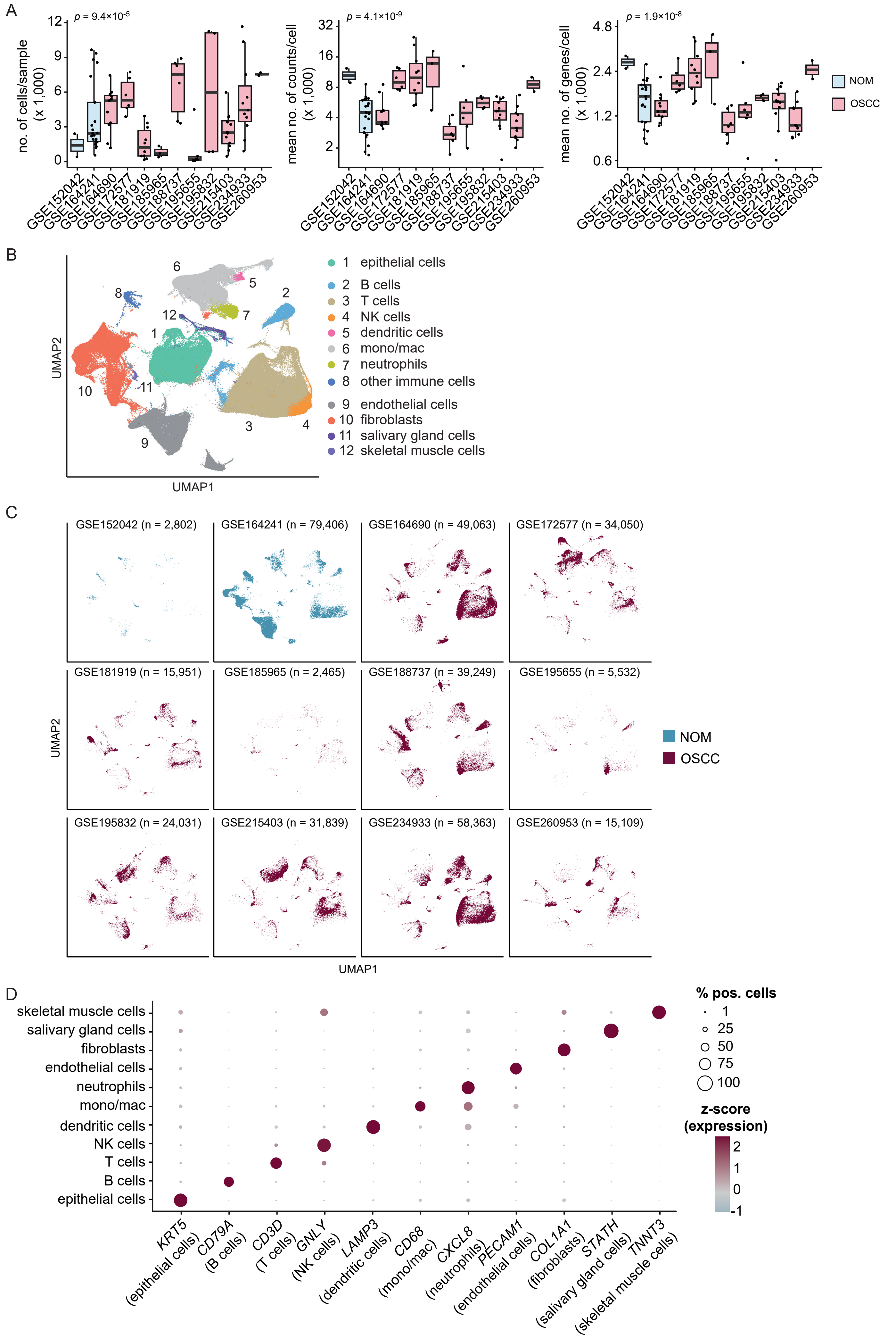


**Supplementary Fig. S1.**

**Legend to Supplementary Figure S1: Integration of publicly available scRNA-seq datasets from OSCC and normal oral mucosa.** (A) Left panel, number of cells per sample; middle panel, sample-level mean number of counts per cell; right panel: sample-level mean number of detected genes per cell. One-way ANOVA. (B,C) Integration of datasets. (B) Integrated UMAP of all OSCC (n = 71) and NOM (n = 23) samples. (C) Representation of cells from each individual dataset (n = 12) in the integrated UMAP. (D) Expression of cell-type specific markers in the respective cells. mono/mac, monocytes/macrophages; NK cells, natural killer cells; NOM, normal oral mucosa; pos. positive.

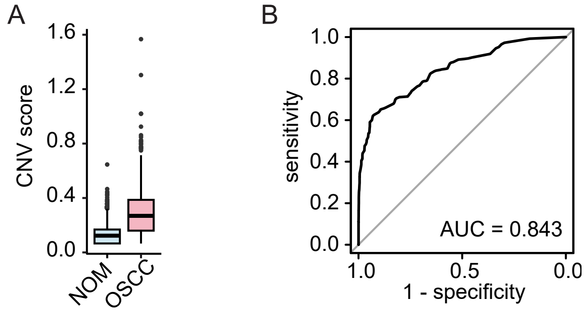


**Supplementary Figure S2: Copy number variations in epithelial cells from OSCC and normal oral mucosa samples.** (A) CNV scores of epithelial cells from normal oral mucosa (NOM; = 5,473) and OSCC (n = 49,093). The CNV score represents the average of the deviation from the diploid state as derived through Infer-CNV for each gene detected in a specific cell. (B) Receiver-operator curve for the CNV score to classify epithelial cells as OSCC or NOM. AUC, area under the curve; CNV, copy number variation; NOM, normal oral mucosa.

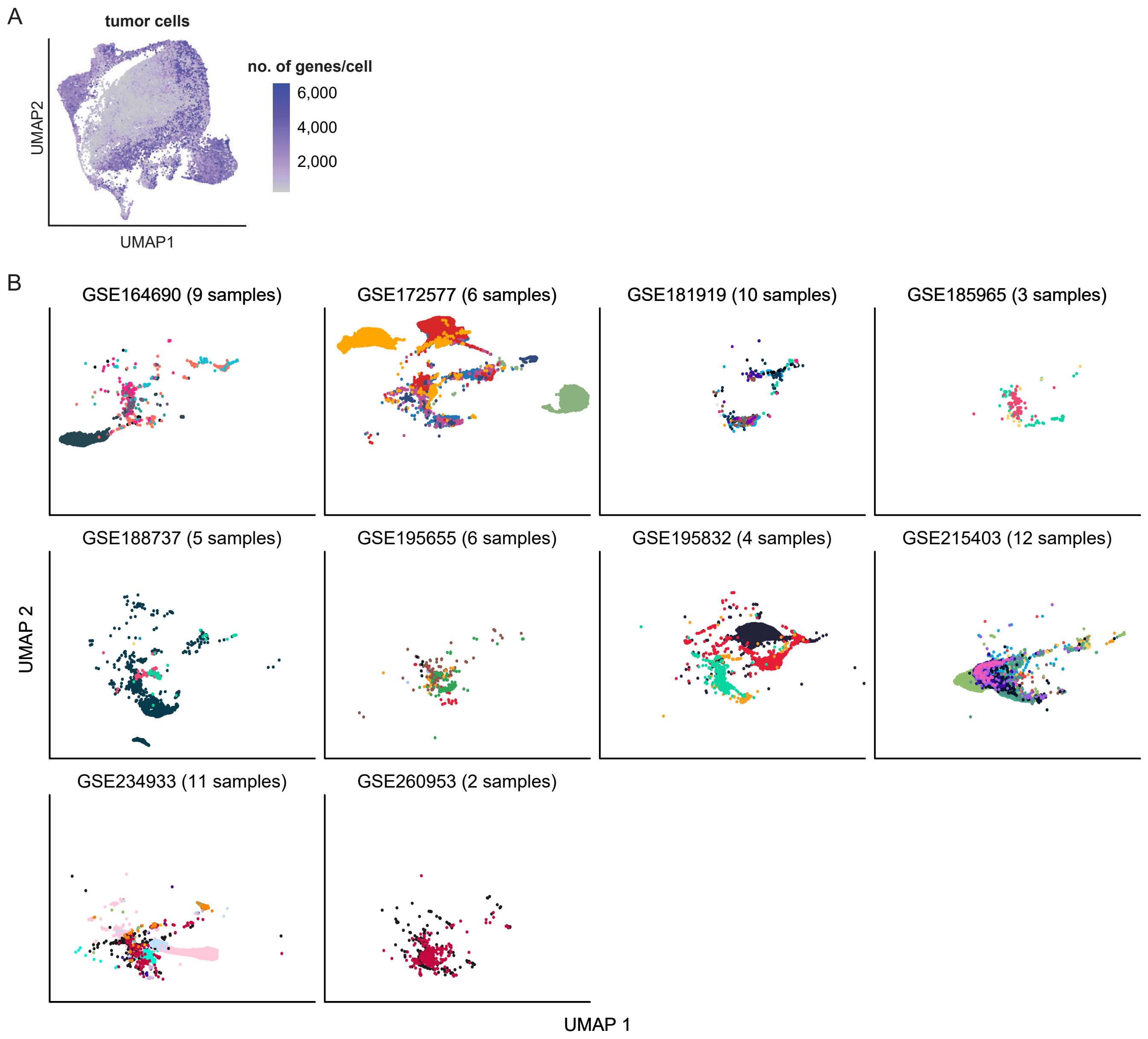


**Supplementary Figure S3: Clustering of tumor cells based on gene expression.** (A) Integrated UMAP of tumor cells (n = 49,093) from OSCC samples (n = 68). (B) Representation of tumor cells from each individual dataset (n = 10) in the integrated UMAP. Each color represents a different sample.

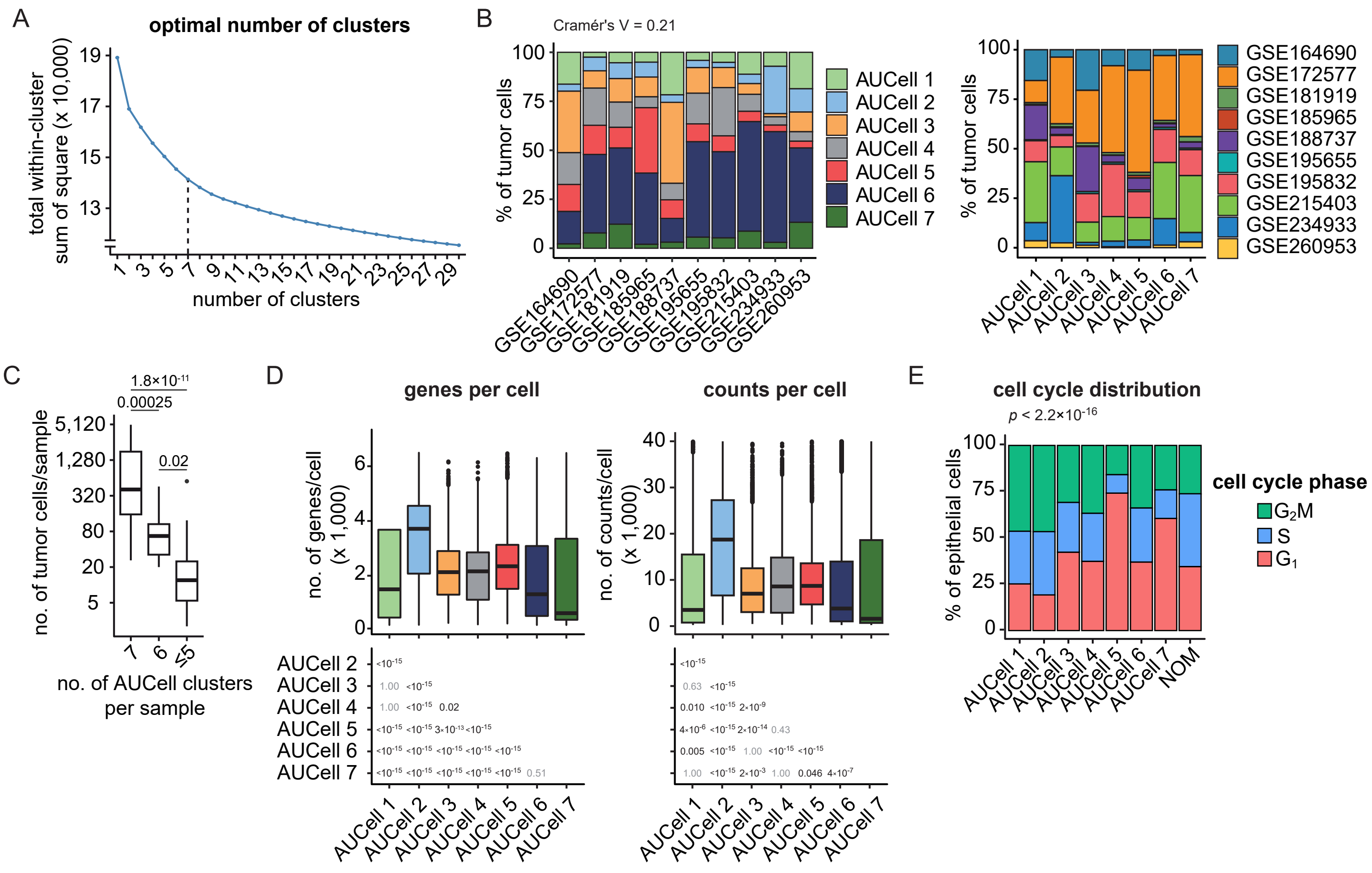


**Supplementary Figure S4: Clustering of tumor cells based on the activation status of the Hallmark Gene Sets from the Molecular Signatures database.** (A) Elbow plot showing that increasing the number of clusters beyond seven does not lead to proportional gains in explained variance. (B) Left panel, relative abundance of tumor cells (n = 49,093) assigned to the different AUCell clusters within each OSCC dataset; right panel, relative abundance of cells from the different OSCC datasets within each AUCell cluster. $p < 2.2 \times 10^{-16}$. Pearson's $X^2$ test. (C) Number of tumor cells in OSCC samples that contained 7 (n = 40), 6 (n = 12), and ≤5 (n = 16) AUCell clusters. One-way ANOVA followed by Bonferroni's post-hoc test. (D) Top left panel, number of detected genes per cell in the different AUCell clusters; top right panel, number of counts per cell in the different AUCell clusters; bottom panels: adjusted p values of all comparisons. One-way ANOVA followed by Bonferroni's post-hoc test. (E) Cell cycle distribution of cells assigned to the different AUCell clusters and of normal oral mucosa epithelial cells, determined using the respective function of Seurat. Pearson's $X^2$ test. NOM, normal oral mucosa.

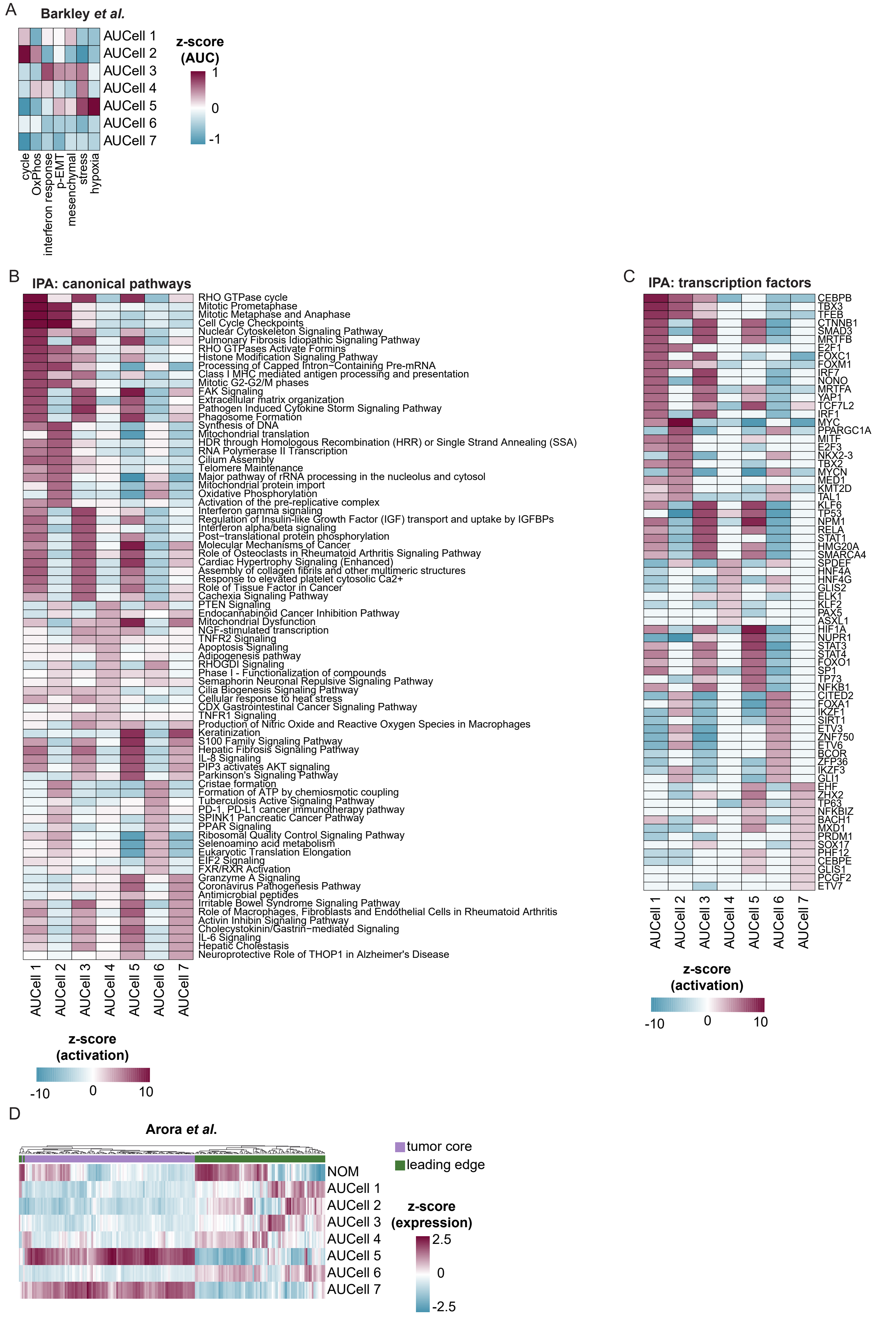


**Supplementary Fig. S5.**

**Legend to Supplementary Figure S5: Gene set enrichment and transcription factor activity in the AUCell clusters.** (A) Enrichment of the tumor-relevant gene sets defined by Barkley *et al*. (37) in the AUCell clusters, determined using *AUCell*. (B) Ingenuity pathway analysis: enrichment of canonical pathways in lists of genes significantly differentially expressed in one vs. all other AUCell clusters. The 15 pathways with the highest z-scores are shown for each cluster, a full list is provided in **Supplementary Table S6**. (C) Transcription factors predicted to explain gene expression differences between the AUCell clusters. Lists of genes significantly differentially expressed in one vs. all other AUCell clusters were analyzed for activity of upstream regulators using IPA software. Results were filtered for transcription factors, and for z-scores >2 and a false discovery rate < 0.05. The 15 transcription factors with the highest z-scores are shown for each cluster; for AUCell 4, only nine transcription factors with increased activity were detected. (B,C) White rectangles indicate pathways and transcription factors with z-scores between 2 and -2 and/or a false discovery rate ≥0.05 (D) Expression of the genes constituting the tumor core and leading edge signatures reported by Arora *et al.* (31) in the AUCell clusters. AUC, area under the curve; IPA, Ingenuity pathway analysis; OxPhos, oxidative phosphorylation; p-EMT, partial epithelial-mesenchymal transition.

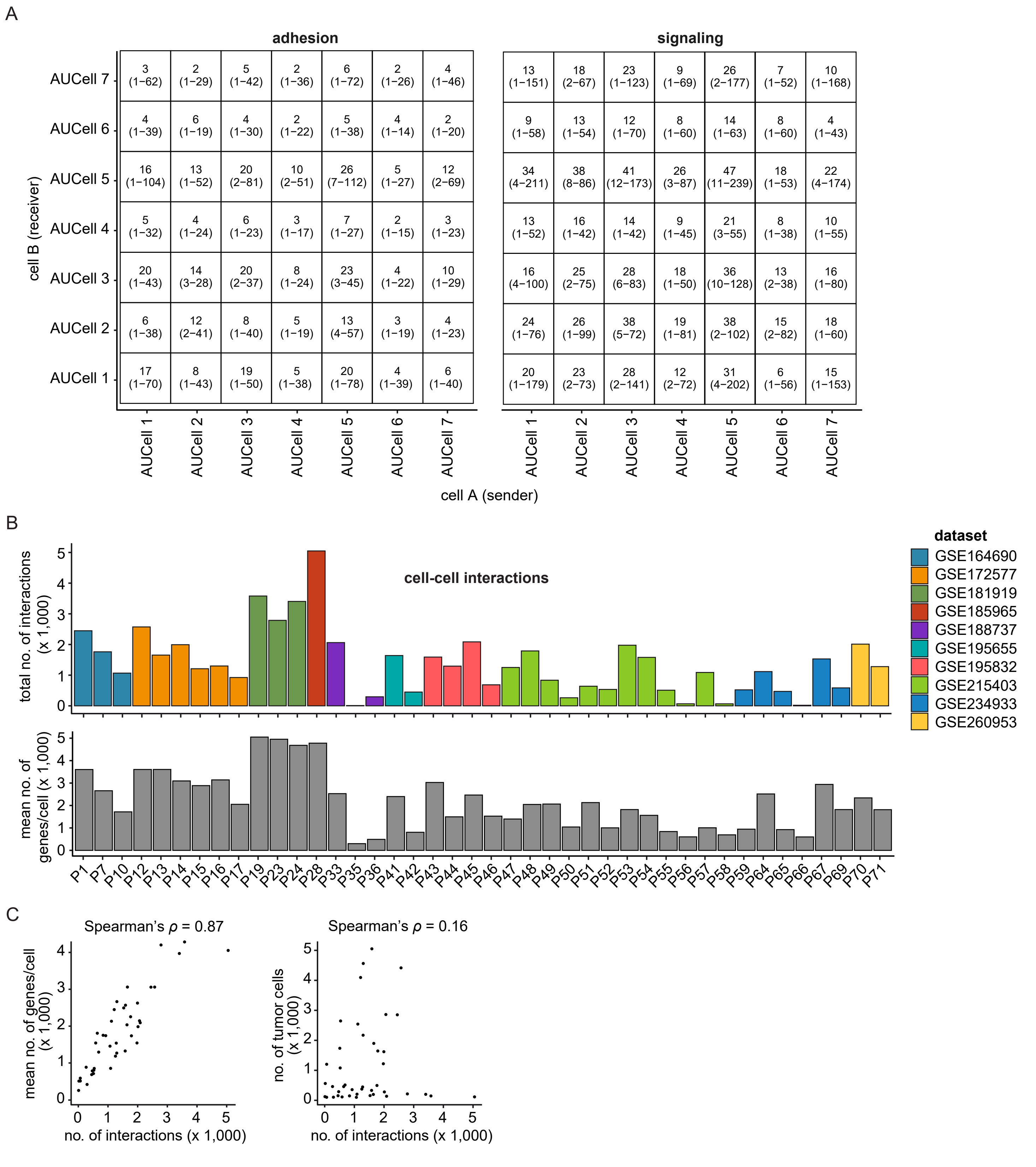


**Supplementary Figure S6: Cell-cell communication within and between AUCell clusters.** (A) Numbers of adhesion and signaling interactions within and between AUCell clusters in individual OSCC samples, determined using CellPhoneDB v5. Mean (range). (B) Top panel, total number of cell-cell interactions for each OSCC sample with ≥100 tumor cells (n = 42). Molecular interactions of the same type but between different pairs of AUCell clusters were counted separately. Bottom panel, sample-level mean number of genes per cell. (C) Correlation between the mean number of genes per cell (left panel), or the number of tumor cells per sample (right panel), and the number of interactions per sample.

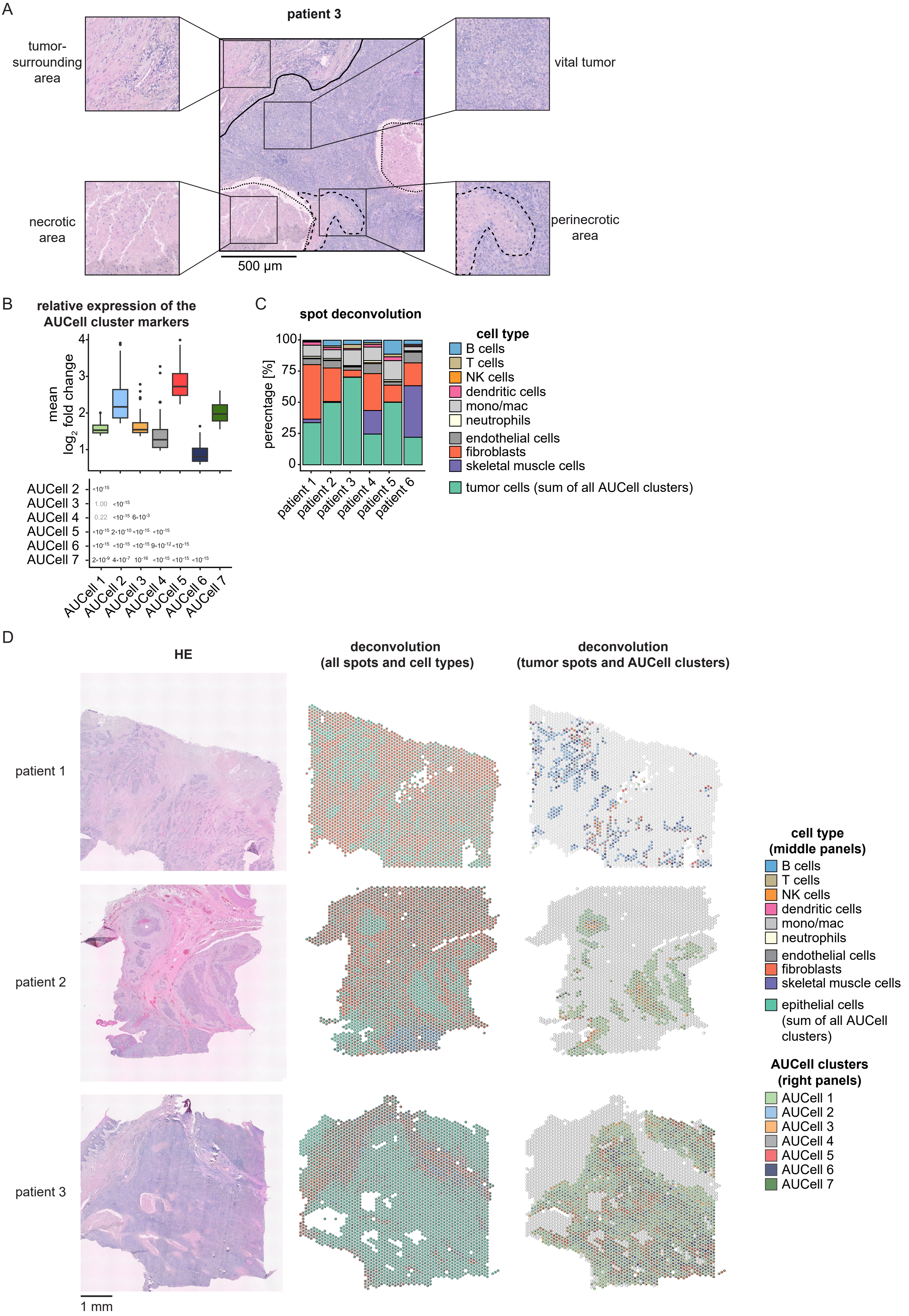


**Supplementary Figure S7.**

D (continued)

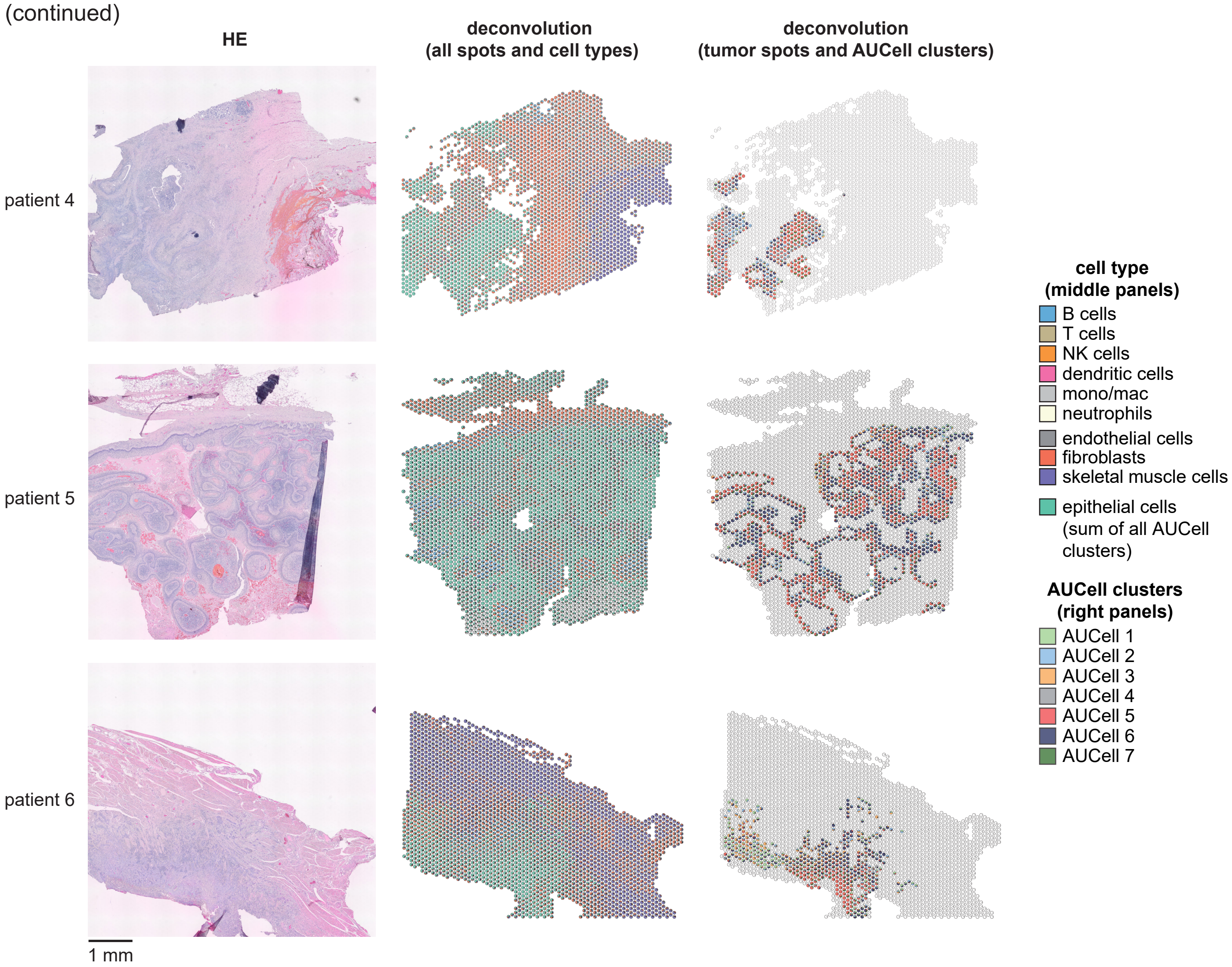


**Supplementary Figure S7: Annotation and deconvolution of spatial transcriptomics data from six OSCC samples.** (A) Histology-based annotation of tumor-surrounding tissue, vital tumor, necrotic and perinecrotic areas, exemplified in the sample from patient 3. Necrotic areas were excluded from further analyses due to low numbers of nuclei and of detected genes. (B-D) Deconvolution of Visium spots (Visium v1; 10x Genomics). The top 71 upregulated marker genes for each cell type and AUCell cluster, derived from the integrated scRNA-seq dataset, were used as reference matrix for *Robust Cell Type Decomposition*. (B) Top panel, mean $\log_2$ fold changes of the top 71 upregulated marker genes for each AUCell cluster relative to all other AUCell clusters; bottom panel, adjusted p values of all comparisons. One-way ANOVA followed by Bonferroni's post-hoc test. (C) Mean proportions of major tumor-resident cell types in spots located in vital tumor and/or perinecrotic areas ($n_{all\ patients}$ = 5,167). AUCell clusters were combined into the category "tumor cells". (D) Left panels, HE images; middle panels, deconvolution of spots covering tumor-surrounding tissue, vital tumor, or perinecrotic areas into the major tumor-resident cell types; right panels, deconvolution of spots covering vital tumor and/or perinecrotic areas into the AUCell clusters. Each spot is represented by a pie chart showing the relative proportions of the cell types indicated in the in-figure legends. Scale bars apply to all panels.

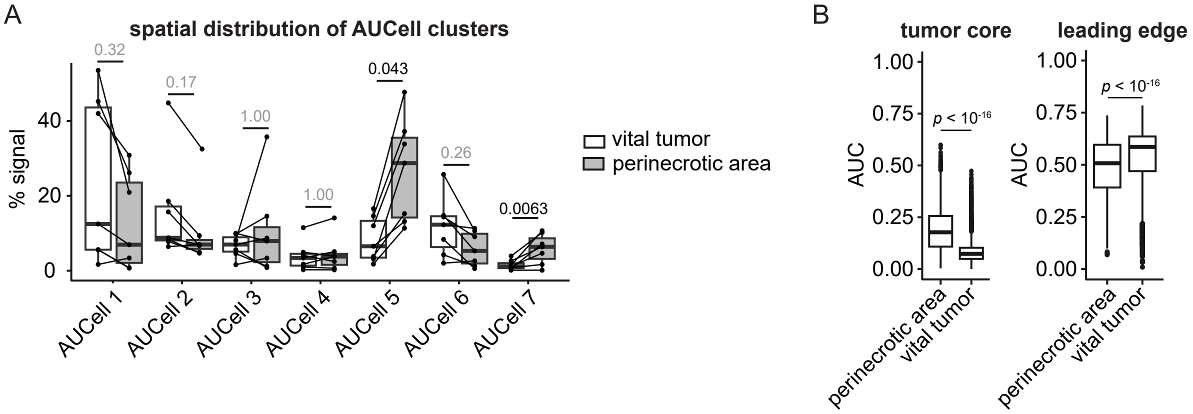


**Supplementary Figure S8: The perinecrotic area is enriched for AUCell 5 and 7 and the tumor core signature.** (A) Proportions of the AUCell clusters in Visium spots in vital tumor vs. perinecrotic areas. Data points represent means from individual patients (n = 6). Two-way ANOVA followed by Bonferroni's post-hoc test. (B) Enrichment of the tumor core (left panel) and leading edge (right panel) signatures in Visium spots in vital tumor (n = 3,917) and perinecrotic (n = 1,250) areas. Two-sided student's *t*-test. AUC, area under the curve.

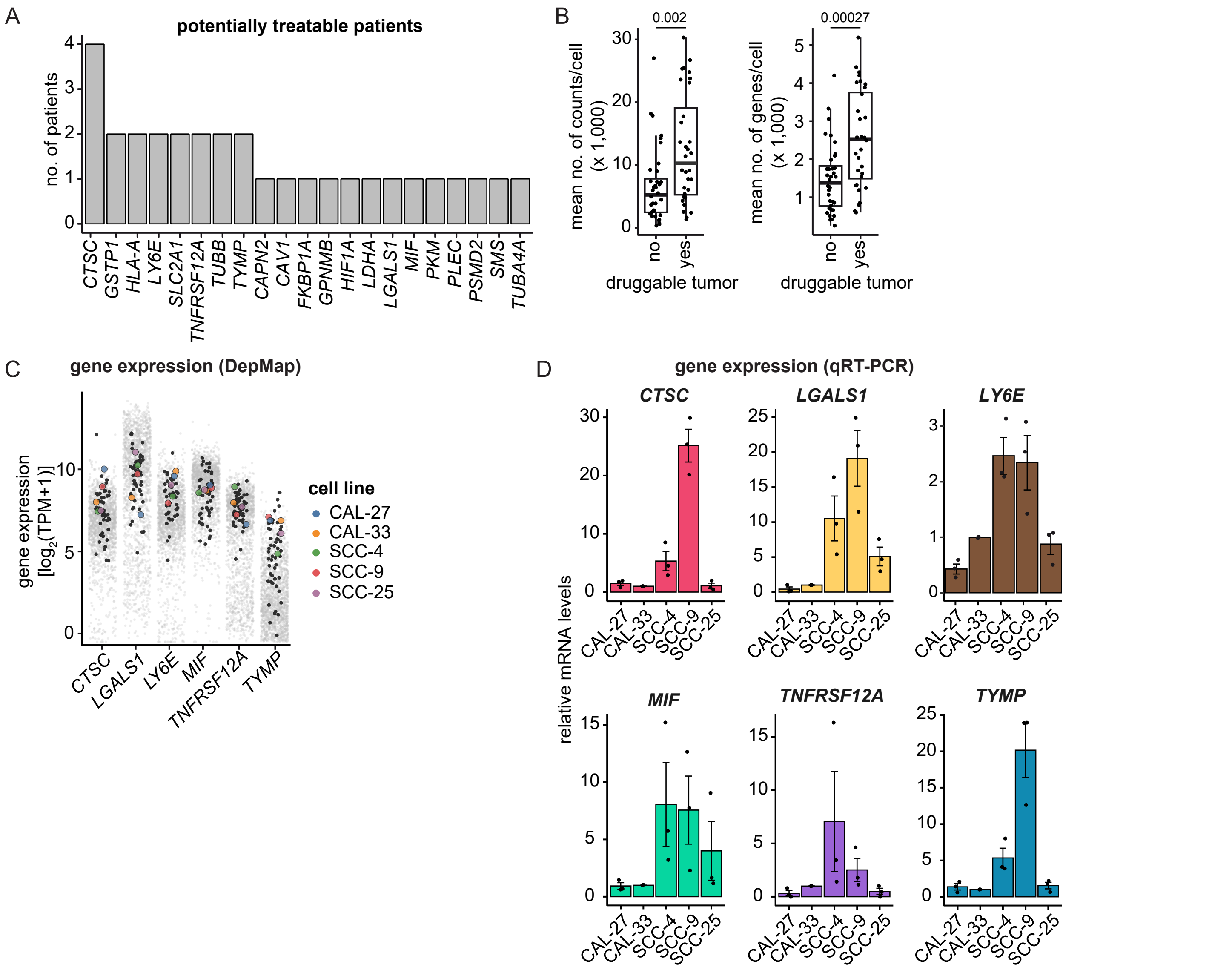


**Supplementary Figure S9: Identification of novel drug combinations for the treatment of OSCC.** (A) Numbers of patients showing overexpression of the indicated drug target genes in >90% of tumor cells. (B) Sample-level mean numbers of counts (left panel) and detected genes (right panel) per cell in OSCC samples defined as “druggable” (n = 32) or “not druggable” (n = 36). Two-sided student’s *t*-test. (C) Expression data for *CTSC*, *LGALS1*, *LY6E*, *MIF*, *TNFRSF12A* and *TMYP* in 1,714 human cell lines from the DepMap database. OSCC cell lines also shown in (D) were highlighted through colored dots and all other OSCC cell lines (n = 54) through black dots. (D) mRNA levels of *CTSC*, *LGALS1*, *LY6E*, *MIF*, *TNFRSF12A* and *TYMP* in five OSCC cell lines as determined by qRT-PCR. Expression levels of the target genes were normalized to those of *B2M* and to CAL-33 using the $\Delta\Delta C_T$ method. Mean ± SEM; n = 3. qRT-PCR quantitative real-time PCR; TPM, transcripts per million.

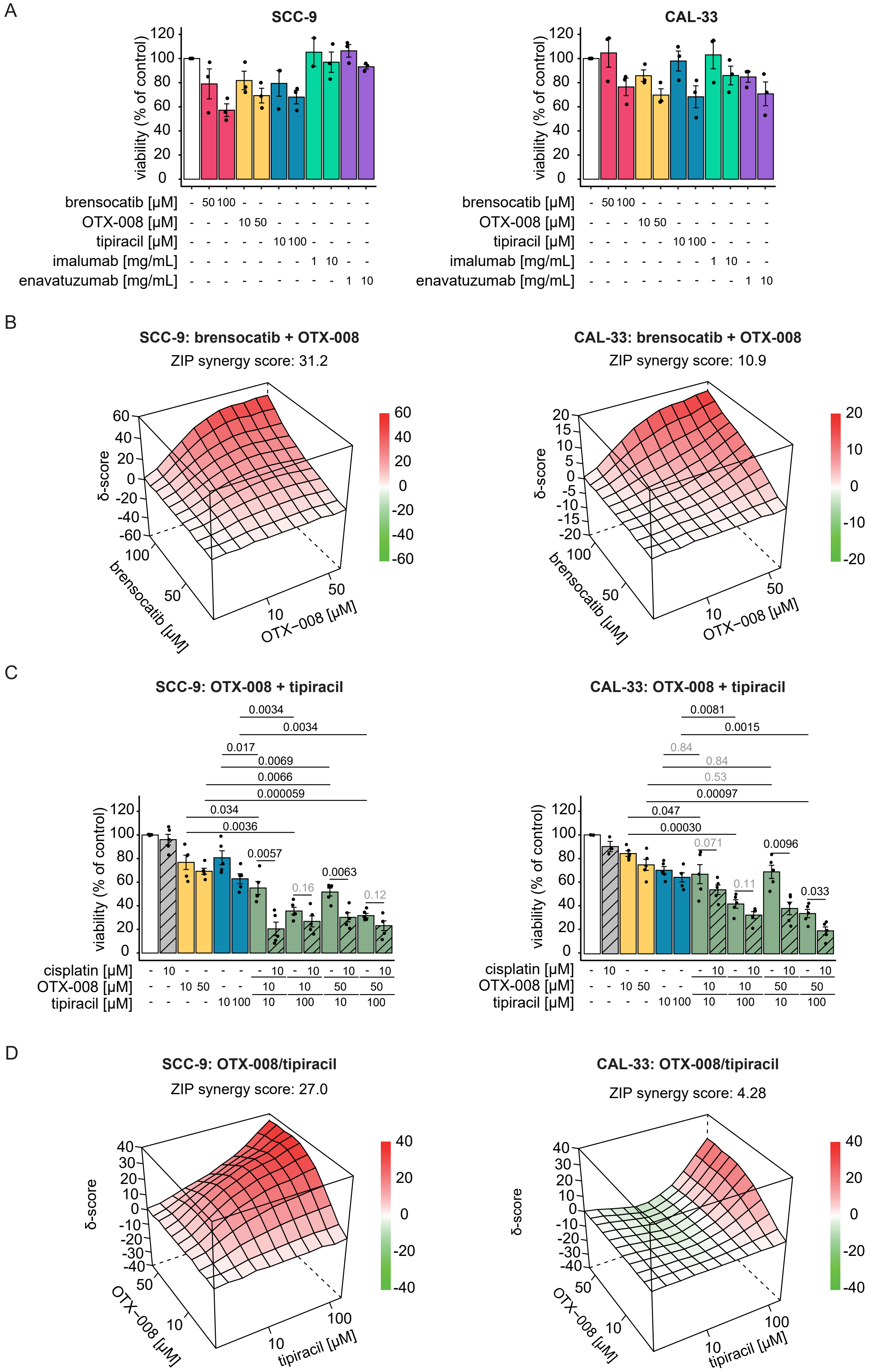


**Supplementary Fig. S10.**

E

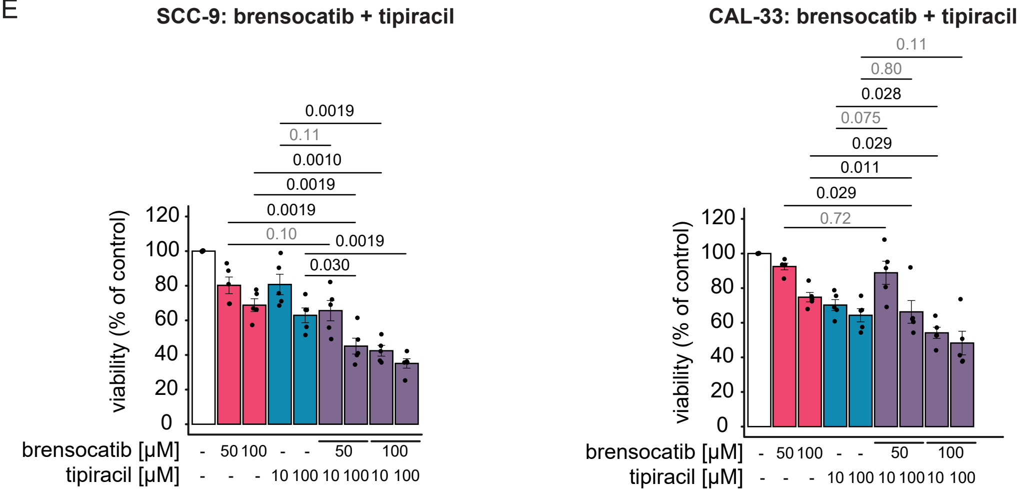


F

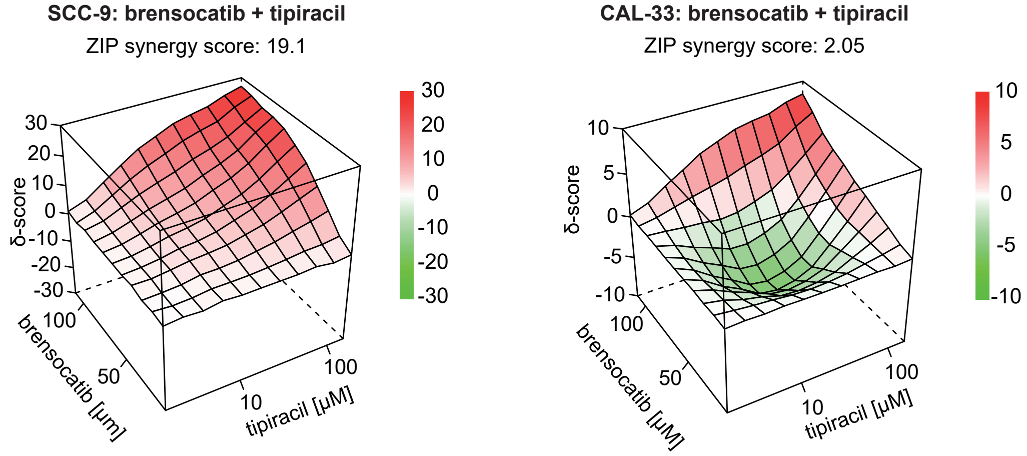


G

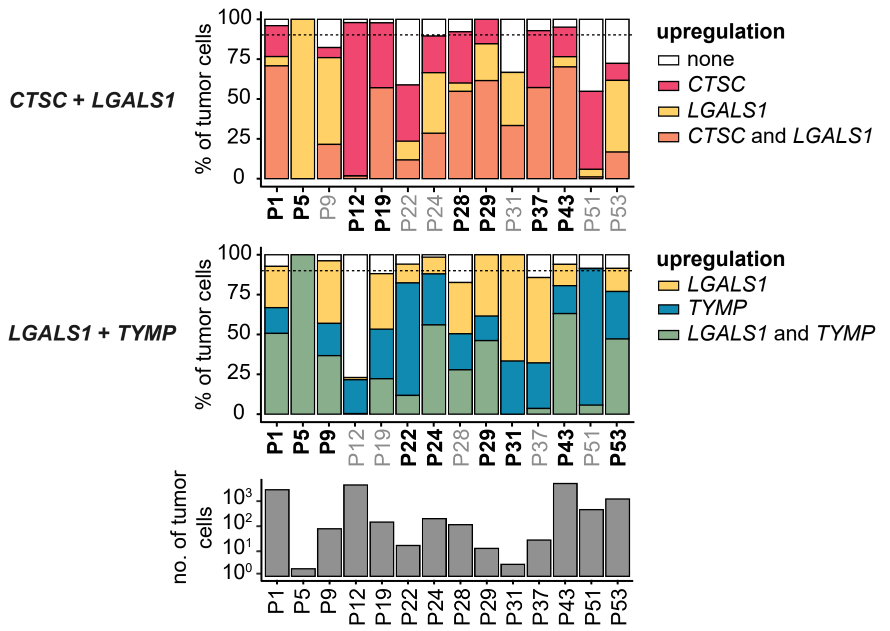


**Supplementary Fig. S10 (continued)**

**Legend to Supplementary Figure S10: Validation of novel drug combinations in tumor spheroids from OSCC cell lines.** (A) Single treatments. Tumor spheroids of the OSCC cell lines SCC-9 (left panel) and CAL-33 (right panel) were treated with the indicated concentrations of the indicated compounds for 5 (SCC-9) or 3 (CAL-33) days and viability was quantified using the CellTiter-Glo 3D Cell Viability Assay. Means ± SEM; n = 3. (B) Synergy scores for the combination of brensocatib with OTX-008. Corresponding viability data are shown in **Figure 5D**. (C-E) Combination treatments. (C) Tumor spheroids of the OSCC cell lines SCC-9 (left panels) and CAL-33 (right panels) were treated with the indicated concentrations of OTX-008, tipiracil and cisplatin individually and in combination for 5 (SCC-9) or 3 (CAL-33) days. (D) Synergy scores for the combination of OTX-008 with tipiracil. (E) Tumor spheroids of the OSCC cell lines SCC-9 (left panels) and CAL-33 (right panels) were treated with the indicated concentrations of brensocatib and tipiracil individually and in combination for 5 (SCC-9) or 3 (CAL-33) days. (F) Synergy scores for the combination of brensocatib with tipiracil. (C,E) Viability was quantified using the CellTiter-Glo 3D Cell Viability Assay. Two-sided student's *t* tests for predefined comparisons, followed by correction for multiple testing according to Bonferroni-Holm. Means ± SEM; n = 5. (F) Proportions of cells overexpressing none, one or both genes of the target combinations *CTSC* plus *LGALS1* (top panel) and *LGALS1* plus *TMYP* (middle panel) in samples defined as "druggable" by one or both combinations, and number of tumor cells per sample (bottom panel).
